# A high-content RNA-based imaging assay reveals integrin beta 1 as a cofactor for cell entry of non-enveloped hepatitis E virus

**DOI:** 10.1101/2023.10.27.564362

**Authors:** Rebecca Fu, Zoé Engels, Jasmin Alara Weihs, Josias Mürle, Mara Klöhn, Daniel Todt, Jungen Hu, Eike Steinmann, Tobias Böttler, Pierre-Yves Lozach, Stanley M. Lemon, Viet Loan Dao Thi

**Affiliations:** Schaller Research Group, Department of Infectious Diseases, Virology, University Hospital Heidelberg, Heidelberg, Germany; Heidelberg Biosciences International Graduate School (HBIGS), Heidelberg, Germany; Department of Molecular and Medical Virology, Ruhr University Bochum, Bochum, Germany; Department of Medicine II, Medical Center – University of Freiburg, Germany; Faculty of Medicine, University of Freiburg, Germany; Center for Integrative Infectious Diseases Research (CIID), University Hospital Heidelberg, Heidelberg, Germany, CellNetworks– Cluster of Excellence, Heidelberg, Germany, Univ. Lyon, INRAE, EPHE, IVPC, Lyon, France; Departments of Medicine and Microbiology & Immunology, The University of North Carolina at Chapel Hill, Chapel Hill, NC, USA; German Centre for Infection Research (DZIF), Partner Site Heidelberg, Heidelberg, Germany

**Keywords:** Hepatitis E virus, cell entry, integrin beta 1, endocytic trafficking, cathepsin

## Abstract

Hepatitis E virus (HEV) is a major cause of acute hepatitis and mainly transmitted faecal-orally. HEV particles in faeces are non-enveloped, while those in the blood possess a cell-derived lipid envelope. Despite being a global health concern, there is limited understanding of the steps in the HEV life cycle, particularly cell entry. A previous study proposed integrin alpha 3 (ITGA3) as a potential host factor for nHEV entry, but the β-integrin partner that co-mediates HEV entry has not been described. To address this knowledge gap and resolve the existing controversies surrounding HEV cell entry, we developed an RNA-FISH-based high-content imaging assay alllowing investigation of the entry pathways of both naked and enveloped HEV particles. Our observations indicate that naked HEV particles interact with the surface receptor integrin beta 1 (ITGB1), which likely facilitates their trafficking through the recycling endosome. In contrast, enveloped HEV particles do not interact with ITGB1 and instead use the classical endocytic pathway via the early endosome. Importantly, both forms of HEV require endosomal acidification and proteolytic cleavage by lysosomal cathepsins, which ultimately results in delivery of the HEV genome to the cytoplasm.

## Introduction

Hepatitis E virus (HEV) is a major cause of acute fulminant hepatitis^1^. HEV infections are usually self-limiting in healthy individuals but can become chronic in immunocompromised patients and cause a high mortality rate in pregnant women and patients with previous liver injury^1^ The virus has a single-stranded, positive RNA genome of 7.2 kb, encoding three open reading frames (ORF1, 2, and 3, Fig. 1A) (reviewed in^2^). ORF1 encodes the non-structural proteins responsible for viral replication, ORF2 the capsid protein, and ORF3 a small phosphoprotein that mediates secretion of viral progenies (Fig. 1a). There are eight HEV genotypes (HEV-1 to 8), of which five (HEV-1 to -4 and HEV-7) are capable of infecting humans^3^. Of note, HEV-3, which is the predominant genotype in developed countries, can infect a broad range of animals, including pigs, deer, and rabbits, and can be transmitted zoonotically to humans. In vitro, HEV-3 can infect a wide range of non-hepatic cell types (reviewed in^4,5^).

**Figure 1:**
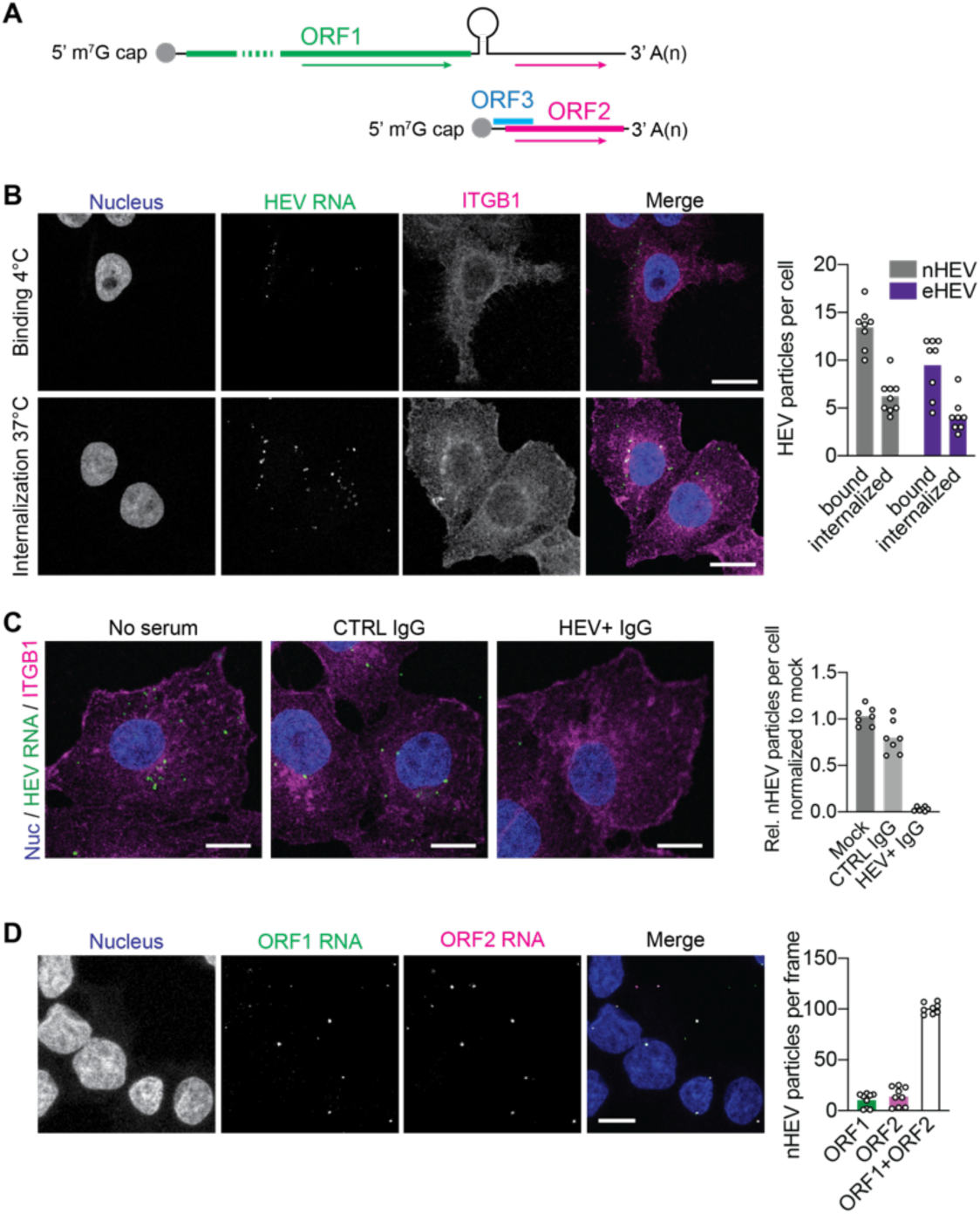
RNA-fluorescence in situ hybridization allows the detection of single HEV particles. (A) Scheme of HEV genome organization showing RNA-fluorescence *in situ* hybridization (RNA-FISH) probes targeting either the full-length genome (green arrow) or the full-length and subgenomic genome (pink arrow). (B) Prechilled hepatoma cells S10-3 were inoculated with nHEV (MOI = 30 GE/cell) or eHEV particles (MOI = 20 GE/cell) and incubated for 2 h or 6 h, respectively, at 4 °C to allow particle binding. The cells were then either fixed or the inoculum removed and shifted to incubation at 37 °C for 6 h to allow HEV particle internalization. After binding or internalization, the cells were fixed and stained with DAPI (blue, nucleus) and against ITGB1 (magenta) and HEV genomes (green) were detected by RNA-FISH (version 2 kit) using the ORF1 probe. Scale bar = 20 μm. (C) nHEV particles were preincubated with convalescent HEV patient serum (1:1000) containing anti-ORF2 antibodies or with non-HEV patient serum as control for 1 h at 37°C. S10-3 cells were inoculated with mock or serum-treated nHEV (MOI = 30 GE/cell) at 37 °C for 6 h. Cells were then fixed and stained as described in (B). Scale bar = 10 μm. (D) S10-3 cells were inoculated with nHEV (MOI = 10 GE/cell) in incubated for 6 h at 37 °C followed by fixation. HEV genomes were detected by RNA-FISH using both ORF1 (green) and ORF2 (magenta) probes. Scale bar = 10 μm. All images represent single slices of confocal images acquired on a Leica SP8 confocal microscope. The detected HEV genomes were quantified using CellProfiler. HEV particles per cell were calculated by dividing the total number of detected HEV genomes by the number of nuclei in an image frame. n = biological replicates from three independent experiments. Images are representatives of n = 7.

HEV is usually transmitted fecal-orally through contaminated food and water^1^. It enters and leaves the human body in its non-enveloped form (nHEV) and circulates in the blood wrapped in a host-derived lipid quasi-envelope (eHEV) acquired during virus budding from cells (reviewed in^6^). eHEV particles in the blood are protected from neutralizing antibodies, while nHEV particles shed in the faeces facilitate transmission outside the host^7^. This directional release is mediated by the polarisation of hepatocytes in vivo, from which apically co-secreted bile acids strip off the envelope as the progeny particles are released into the bile^8^. The quasi-envelope confers the particles a lower buoyant density, making them approximately ten times less infectious than the naked virions^8–10^. In this respect, HEV is very similar to hepatitis A virus (HAV), which is also enterically transmitted and occurs in both, naked and quasi-enveloped forms^11^.

Despite growing interest, fundamental steps of the HEV life cycle, including the cell entry pathways of both nHEV and eHEV particles are poorly understood. Several host factors, such as heparan sulfate proteoglycans, glucose-regulated protein 78, and asialoglycoprotein receptor have been proposed as nHEV attachment factors (reviewed in^12^). While an HEV entry receptor has not yet been identified, epidermal growth factor receptor (EGFR) has recently been shown to modulate HEV entry in human hepatocytes^13^. Additionally, a very recent study showed that the T cell immunoglobulin mucin domain-1 (TIM-1) receptor promotes infection of eHEV particles through binding to phosphatidylserines which are present on their quasi-envelope^14^.

Shiota and colleagues have previously proposed integrin α3 (ITGA3) as another entry factor for HEV^15^. The authors demonstrated a strong correlation between ITGA3 expression and nHEV permissiveness as well as a direct interaction between nHEV particles and the ITGA3 ectodomain. Integrins are cell surface receptors involved in various cellular processes including cell adhesion, cell migration, signal transduction, proliferation, and apoptosis (reviewed in^16^). They are transmembrane proteins that always function as αβ dimers. Heterodimerisation of eighteen α-integrin and eight β-integrin subunits result in the assembly of 24 distinct integrin receptors^16^. However, the β-integrin partner that co-mediates HEV entry has not been described yet.

It has recently been shown that the two types of HAV particles enter the cell via similar endocytic pathways, with transfer of the HAV genome to the cytoplasm dependent upon ganglioside receptors in the late endolysosome^17^. However, studies on the involvement of endosomal trafficking of HEV particles have been contradictory. Following potential receptor interactions, Yin and colleagues found that both eHEV and nHEV particles are internalised via clathrin-dependent endocytosis^9^. However, this study also proposed that eHEV entry requires the small GTPases Rab5 and Rab7, whereas nHEV entry did not appear to rely on the endocytic pathway. In contrast, the Holla *et al.* study based on virus-like particles (VLPs) showed that HEV VLPs traffic to Rab5-positive compartments en route to acidic lysosomal compartments where they are degraded^18^.

The absence of suitable assays likely contributes to the current lack of understanding of the molecular steps of HEV cell entry. Unlike enveloped viruses, HEV particles lack glycoproteins and cannot be studied using viral pseudotypes. The capsid itself is very compact and fusion with fluorescent tags is likely to affect both capsid assembly and receptor interactions during entry. To address this gap in knowledge and controversies surrounding HEV cell entry, we developed a high-content imaging approach to visualise and study the entry pathways of both naked and quasi-enveloped HEV particles. Our studies reveal a critical role for the cell surface receptor integrin β1 (ITGB1) in determining the endocytic trafficking pathways of nHEV, but not eHEV particles.

## Materials and Methods

### Cell culture

The human hepatoma cell lines S10-3 (a kind gift from Suzanne Emerson, NIH) and HepG2.C3A and their derivatives were cultured in Dulbeccos’s Modified Eagle Medium (DMEM, Gibco) + GlutaMAX-I supplemented with 10% fetal bovine serum (FBS, Capricorn), 1% penicillin/streptomycin (P/S) (Gibco) here referred to as complete DMEM (cDMEM). Cell lines were validated by phenotypic screening and confirmed to be mycoplasma-free using a PCR detection kit (Abcam). All cells were maintained at 37 °C in a 95% humidity and 5% CO_2_ atmosphere.

### Production and purification of nonenveloped and enveloped HEV particles

nHEV and eHEV particles were harvested from S10-3 cells electroporated with *in vitro* transcribed HEV GT3 Kernow C1 P6 (GenBank accession number: JQ679013.1) RNA, 7 days post-electroporation. In brief, nHEV particles were collected from the cell lysate through four freeze-thaw cycles, and eHEV particles were harvested from the filtered cell culture supernatant. The HEV particles were then concentrated by layering on top of a 20% w/v sucrose cushion and ultracentrifuged at 28000 rpm using a SW 32 Ti Swinging-Bucket Rotor for 3 h at 4 °C. The resulting pellet was resuspended in PBS and further purified with help of a continuous gradient, which was prepared by layering 2.5 ml of 60%, 40%, 25%, and 15% Opti-prep^TM^ (Sigma) (v/v) each in a Thinwall Ultra-Clear Tube (Beckman). The gradient was placed horizontally for 1 h to allow mixing of the gradient and kept overnight at 4 °C. The next day, concentrated virus was layered on top of the gradient and centrifuged at 32000 rpm for 16 h at 4 °C using a SW 40 Ti Swinging-Bucket Rotor. Twelve individual 1 ml gradient fractions were manually collected. The refractive index of each fraction was measured using a digital handheld refractometer (DR201-95, Krüss). Fractions with a density ranging from 1.05 to 1.15 g/cm^3^ were pooled for eHEV and from 1.2 to 1.25 g/cm^3^ for nHEV infection experiments. Prior to infection, the gradient-purified virus was subjected to buffer exchange using the Pur-A-Lyzer™ Maxi Dialysis Kit (20k MWCO) (Sigma) at 4°C overnight to replace the iodixanol with PBS.

### HEV infection and neutralization assays

3×10^4^ S10-3 cells were seeded in a well of a 48-well plate and infected with nHEV or eHEV in MEM (Gibco) supplemented with 10% FBS and 1% P/S, the next day. The inoculum was removed after 8 h and replaced with cDMEM. 5 days post-infection, the cells were fixed in 4% PFA (Electron Microscopy Sciences) for 15 min and permeabilized with methanol at −20 °C for 20 min. The cells were then blocked with 10% goat serum for 1 h at room temperature, immunostained with an ORF2 antibody (1E6 1:1000, Millipore) at 4 °C overnight followed by anti-mouse Alexa-594 (1:1000, Thermo Fisher) staining for 1 h at room temperature. All infection assays were carried out at 37°C. Images of entire infected wells were taken with a Zeiss CellDiscoverer 7 microscope with a 10x objective and the number of FFU was counted manually. For neutralization, 4×10^4^ S10-3 cells were seeded on a 10 mm coverslip in a well of a 48-well plate. Gradient-purified nHEV was neutralized with convalescent patient-serum or HEV-negative patient serum at 1:1000 for 1 h at 37°C. The cells were then inoculated with serum-treated or untreated nHEV for 6 h at 37°C. The use of the patient-serum was approved by the Ethics Committee of the Albert-Ludwigs-University Freiburg (474/14, 201/17, 486/19), and written informed consent was obtained from all blood donors before enrolment in the study.

### Generation of ITGB1 knockout cell lines

S10-3 cells were transfected with the pSpCas9(BB)-2A-GFP (Addgene #48138) plasmid encoding the Cas 9 protein fused to GFP by the 2A peptide, and the sgRNA sequence (AGAATTTCAGCCTGTTTACA) directed against exon 7 (out of 16). GFP-positive clones were FACS-sorted 48 h after transfection, and single clones were cultured in conditioned media from confluent S10-3 cell cultures supplemented with 20% FBS.

### Ectopic ITGB1 expression and selection

The ITGB1 gene was cloned into the lentiviral expression plasmid pWPI (Addgene #12254) and the Rab 5-EGFP, Rab 7-EGFP, Rab 11-EGFP, and EGFP-LAMP1 genes into the doxycycline-inducible lentiviral expression plasmid pTRIPZ (Thermo Fisher). Lentiviruses were produced by transfecting HEK293T cells with plasmids encoding VSV-G, HIV gag/pol proteins, and the transgene using the JetPRIME reagent (Polyplus) according to the manufacturer’s protocol. Lentiviruses were harvested 48 h post-transfection. S10-3 ITGB1 KO cells were transduced with pWPI-ITGB1 and selected in cDMEM supplemented with 400 μg/ml G418 (Invivogen). S10-3 WT cells were transduced with pTRIPZ-Rab5/7/11-GFP or EGFP-LAMP1 and selected in cDMEM supplemented with 2 μg/ml puromycin (Invivogen).

### siRNA reverse transfection

SMARTpool siRNAs (Dharmacon) were individually added to each well of a 48-well plate containing 25 μl OptiMEM and 1 μl Lipofectamine™ RNAiMAX Transfection Reagent (Thermo Fisher) at a final concentration of 100 nM. After 5 mins incubation at RT, 5×10^4^ S10-3 cells were added to each well in 250 μl culture medium. 48 h later, the cells were inoculated with HEV and harvested for western blot analysis.

### In situ labeling of viral RNA and immunoflourescence staining

Infected S10-3 cells seeded onto coverslips were fixed in 4% PFA and permeabilized in 0.1% Trion-X100 (Sigma). For co-detection of RNA and capsid, RNAscope® Fluorescent Multiplex Kit version 1 (ACDBio) was used according to the manufacturer’s protocol. The positive strand of HEV RNA was targeted by the ORF1 probe (ACDBio, Cat No. 579831) or ORF2 probe (ACDBio, Cat No. 586651). Subsequently, cells were blocked in 5% goat serum followed by immunostaining with 1E6 (Millipore, 1:400) or ITGB1 antibodies (Santa Cruz, 1:100) at room temperature for 1 h. The respective Alex-conjugated secondary antibodies were used at 1:1000 diluted in 5% goat serum and incubated at room temperature for 1 h. Finally, the coverslips were mounted using the ProLong™ Glass Antifade Mountant (Thermo Fisher Scientific) and cured for at least 24 h in the dark. RNAscope® Fluorescent Multiplex Kit version 2 (ACDBio) was used for detection of only RNA. Cells were fixed and permeabilized as described above, followed by H2O2 treatment for 10 min at RT before proceeding with the RNAscope protocol.

### Western blot

Cells were lysed in RIPA lysis buffer (Thermo Fisher) on ice for 30 min and the protein concentration was determined using the Pierce™ BCA protein assay kits (Thermo Fisher) to ensure equal loading. A total of 20 μg of proteins were mixed with 6x SDS loading dye containing 10% 2-mercaptoethanol (VWR Life Sciences) and boiled at 100 °C for 10 min before loading. Proteins were then transferred to polyvinylidene difluoride (PVDF, G-Biosciences) membranes by wet blotting using standard methods. The membranes were blocked with 5% milk/0.1% Tween-20 in PBS (PBS-T). Antigens were stained with the indicated antibodies in 5% milk: mouse α-ITGB1 1:500 (Santa Cruz); mouse α-actin (Abcam) 1:4000; mouse α-FAK (Santa Cruz) 1:500; rabbit α-Rab 7 (Abcam) 1:1000; rabbit α-Rab 11 (Abcam 1:1000); and goat α-Rab 5 (antibodies-online) 1:1000, followed by staining with corresponding secondary antibodies conjugated with HRP. Membranes were imaged with Pierce™ Enhanced Chemiluminescence (ECL) Western Blotting Substrate and images were acquired using ChemoStar Touch ECL & Fluorescence Imager (Intas).

### Confocal microscopy and image analysis

Multichannel z-series with a z-spacing of 10 μm, or single slice confocal images were acquired using a Leica LIGHTNING SP8 or a Zeiss Airyscan LSM900 confocal microscope, as indicated in the figure legend. A 63× oil immersion objective was used for all images. For quantification of HEV genomes per cell during entry and percentages of HEV genomes associated with capsid, maximum projections of full z-series were used. The genomes per cell were estimated by dividing the total number of detected genomes by the number of nuclei in a frame. Images were processed using the Zen software and inspected manually before quantifications using CellProfiler. For colocalization studies, maximum projections of 3-4 z-slices were shown. Intensity plot profiles were generated using the ImageJ software.

### Statistical analysis

Graphs and statistical analyses were performed using GraphPad PRISM 8. In all figures where p-values were calculated, the corresponding statistical test is listed in the figure legend.

## Results

### In situ RNAscope hybridization allows detection of single HEV particles

Previous studies of HEV cell entry have relied on either the use of VLPs^18^ or the quantification of infected cells^9^. However, HEV VLPs are derived from self-assembled truncated recombinant ORF2 proteins (reviewed in^19^) and lack the critical C-terminal protruding “P” domain that is likely responsible for virus binding to cell receptors. In addition, the study of cell entry requires separation from the other steps of the viral life cycle. In the absence of a traditional lipid envelope containing viral glycoproteins, we developed a high-content imaging assay to visualise and study incoming HEV particles based on the detection of single HEV genomes by RNA fluorescence in situ hybridisation (RNA-FISH) (Fig. 1A).

In conventional non-polarised HEV-replicating cell culture systems, naked HEV particles can be recovered from cell lysates, whereas quasi-enveloped particles are released into the supernatant. We density-gradient purified naked (nHEV, 1.25 g/cm^3^) and quasi-enveloped (eHEV, 1.12 g/cm^3^) HEV GT3 Kernow C1 P6 particles (Suppl. Fig. 1A) and allowed them to bind and/or internalise into prechilled S10-3 hepatoma cells before fixing the cells and staining bound genomes using specific RNA-FISH probes targeting ORF1 (Fig. 1A). As shown in Fig. 1B, we detected and quantified individual HEV genomes either at the edges of the cells (bound) or translocated into the interior of the cells (internalised). We then calculated the amount of HEV particles by dividing the number of HEV genomes detected by the number of nuclei in an image frame (Fig. 1A, right panel).

To ensure that the HEV genomes detected were capsid-dependent, we pre-treated nHEV particles with convalescent HEV patient serum containing anti-ORF2 antibodies^20^ or with non-HEV patient serum as a control. While the control serum had little effect on the number of detected internalised HEV genomes, we observed an almost 100% reduction in HEV genomes after neutralisation with patient serum compared to untreated virus (Fig. 1C). In addition, to rule out the potential detection of free HEV genomes in the virus preparation, we pretreated nHEV particles with RNAse A prior to inoculation into S10-3 cells. As shown in Suppl. Fig. 2, the RNAse treatment did not reduce the amount of HEV genomes, thus excluding detection of free HEV genomes in our assay. Taken together, these observations suggest that we successfully detected single viral particles by RNA-FISH.

To ensure that viral particles packaged full-length HEV genomes, rather than defective truncated or subgenomic genomes, we used RNA-FISH probes targeting both ORF1 (green probe) and ORF2 (magenta probe) simultaneously (Fig. 1A). As shown in Fig. 1D, multiplexing of both probes revealed that the majority of detected particles contained the full-length HEV genomes, while only a fraction of particles contained either truncated (ORF1 only) or potentially subgenomic (ORF2 only) genomes. Taken together, our results demonstrate that the RNA-FISH assay can be used to visualise infectious HEV particles during cell entry.

### ITGB1 is a co-host factor of nHEV but not eHEV cell entry

A previous study proposed ITGA3 as a potential entry factor for nHEV^15^. Integrins are obligate heterodimers and ITGA3 appears to only dimerise with ITGB1^15,16^. ITGB1 is ubiquitously expressed, including in primary human hepatocytes (Suppl. Fig. 3A). Since ITGB1 has been shown to mediate entry of many viruses^21–24^, we wanted to investigate whether ITGB1 is also an entry factor for HEV. Using CRISPR-Cas9, we generated ITGB1 knockout (KO) S10-3 cells and selected two clones for our studies (Fig. 2A). We then infected these KO clones with nHEV and eHEV. Throughout our study, we used a higher MOI for eHEV in order to achieve equivalent numbers of FFUs, since eHEV particles are less infectious and have a lower specific infectivity than nHEV particles^7,8^ (Suppl. Fig. 4).

**Figure 2:**
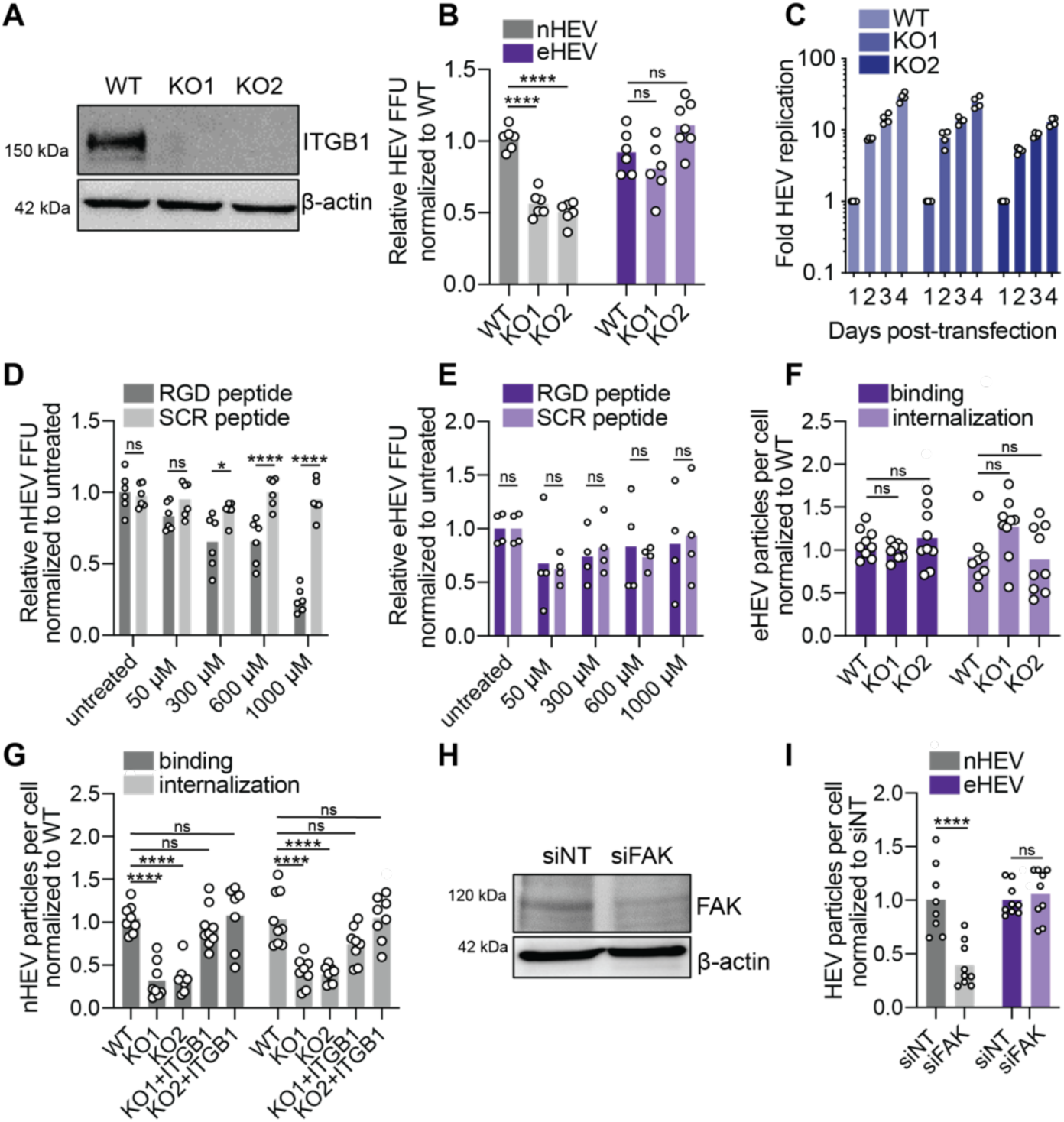
Integrin beta 1 is a co-host factor for nHEV but not for eHEV cell entry. (A) Western blot analysis of S10-3 ITGB1 WT and KO cell lysates. (B) S10-3 ITGB1 WT and KO cells were infected with nHEV (MOI = 0.1 GE/cell) or eHEV particles (MOI = 5 GE/cell). Infectivity was assessed by staining against the capsid protein ORF2 and quantifying FFUs 5 days post-infection. (C) S10-3 ITGB1 WT and KO cells were electroporated with an HEV-GLuc subgenomic replicon and HEV replication was quantified by measuring luciferase activity in the supernatant during days 1 to 4 post-electroporation. (D) and (E) S10-3 WT cells were infected with nHEV (MOI = 0.1 GE/cell) or eHEV particles (MOI = 5 GE/cell) in the presence of indicated concentrations of a RGD-containing peptide or a scrambled (SCR) control peptide. Infectivity was assessed by staining against the capsid protein ORF2 and quantifying FFUs 5 days post-infection. (F) Prechilled S10-3 ITGB1 WT and KO cells were inoculated with eHEV particles (MOI = 20) and incubated for 12 h at 4 °C to allow binding. For internalization, the inoculum was removed after binding and the cells were shifted to 37 °C for 24 h. After binding or internalization, cells were fixed and HEV genomes detected by RNA-FISH (version 2 kit) using the ORF1 probe. The number of bound and internalized eHEV particles were quantified using CellProfiler and normalized to WT cells. (G) Prechilled S10-3 ITGB1 WT and KO cells with and without ectopic ITGB1 expression were inoculated with nHEV particles (MOI = 30 GE/cell) and incubated for 2 h at 4 °C to allow binding. For internalization, the inoculum was removed after binding and the cells were shifted to 37 °C for 6 h. After binding or internalization, cells were fixed and analysed as described in (F). The number of bound and internalized nHEV particles were quantified using CellProfiler and normalized to WT cells. (H) Western blot analysis of lysates harvested from S10-3 cells 48 h post-transfection with 100 nM on-target pool siRNAs directed against the FAK gene (siFAK) or a non-target control (siNT). (I) siFAK- and siNT-transfected S10-3 cells were inoculated with nHEV (MOI = 30 GE/cell) or eHEV particles (MOI = 20 GE/cell), 48 h post-transfection. After incubating the cells for 8 h at 37 °C, they were fixed and analysed as described in (F). For panels (B), (D) and (E), images of entire infected wells were taken with a Zeiss CellDiscoverer 7 microscope and the number of FFU was counted manually. For panels (F), (G), and (I), the images were taken on a Zeiss Airyscan LSM900 confocal microscope. Maximum projections of full z-series with a thickness of 10 μm were used for quantification of RNA and capsid particles with CellProfiler. n = biological replicates from three independent experiments. Statistical analysis was performed by one-way ANOVA **: *p* <0.01; ***: *p*<0.001; ****: p<0.0001; ns, non-significant.

nHEV infection of ITGB1 KO cells was greatly reduced compared to wild-type (WT) cells, as assessed by counting HEV foci forming units (FFUs) 7 days post-infection (Fig. 2B). In contrast, eHEV infection appeared to be unaffected by the absence of ITGB1. As the reduction in nHEV infection could be the result of either an effect on virus entry or genome replication, we next investigated the effect of ITGB1 KO on HEV replication. We electroporated ITGB1 KO and WT cells with a subgenomic Gaussia luciferase (GLuc) reporter HEV replicon and quantified secreted GLuc as a surrogate for HEV replication. Compared to WT cells, HEV replication was not significantly impaired in ITGB1 KO cells (Fig. 2C), suggesting a role for ITGB1 in HEV cell entry.

To complement these results, we blocked ITGB1 with an arginine-glycine-aspartic acid (RGD) peptide, which is a common motif on many integrin ligands. A scrambled RGD peptide was used as a control. Application of the RGD, but not the control peptide, during entry resulted in a dose-dependent inhibition of nHEV but not eHEV infection (Fig. 2D & E). We next sought to validate the role of ITGB1 in HEV entry by studying incoming HEV particles based on HEV genome detection by RNA-FISH. We inoculated WT and KO cells with nHEV and eHEV on ice, followed by inoculum removal and internalisation at 37°C. Consistent with the lack of inhibition of eHEV infection by the RGD peptide (Fig. 2E), we observed no differences in eHEV binding and internalisation in WT or KO cells (Fig. 2F). In contrast, we found that the number of bound and internalised nHEV particles was significantly lower in KO cells than in WT cells (Fig. 2G). To confirm that the reduction in nHEV entry in KO cells was specifically due to the absence of ITGB1, we ectopically rescued the expression of ITGB1 in KO cells (Suppl. Fig. 5A) and found that ectopic expression restored nHEV binding and internalisation similar to WT levels (Fig. 2G).

To further support the role of ITGB1 in HEV entry, we transfected S10-3 cells with small interfering RNA targeting ITGB1 (siITGB1) or non-targeting siRNA (siNT) 48 h prior to HEV inoculation (Suppl Fig. 5B). Consistent with our results in the KO cells, we found that the ITGB1 knockdown significantly reduced nHEV but not eHEV binding and internalisation (Suppl Fig. 5C). Focal adhesion kinase (FAK) is a key factor in integrin activation and internalisation downstream of receptor engagement^25^. All β1 integrins are capable of activating FAK, leading to the recruitment of structural proteins and cytoskeletal rearrangements^26^. Since ITGB1 appeared to play a role in nHEV binding and internalisation, we tested whether FAK is also involved in nHEV entry. As shown in Fig. 2I, knockdown of FAK (Fig. 2H) resulted in a significant reduction of nHEV but not eHEV binding and internalisation. Taken together, our results show that ITGB1 plays a role in nHEV, but not eHEV entry.

### Co-detection of capsid and RNA allows analysis of the dynamics of HEV entry

Successful viral entry requires the release of the viral genome into the cytoplasm where the incoming genome can be translated into non-structural proteins to initiate genome replication. This process requires dissociation of the viral genome from the capsid. To study the kinetics of HEV uncoating, we combined viral genome detection by RNA-FISH with capsid staining using a specific anti-ORF2 antibody. First, we imaged cell-free nHEV particles and observed that the majority of detected HEV genomes overlapped with detected capsids, while only a fraction of capsids appeared to be devoid of HEV genomes (Fig. 3A). We then imaged HEV particles after binding to cells (0 h) and 24 h after internalisation (Fig. 3B). Initially, we observed that all detected HEV genomes colocalised with the capsids, whereas 24 h later, most HEV genomes no longer colocalised with the capsids, indicating successful uncoating and delivery of the genomes to the cytoplasm (Fig. 3B).

**Figure 3.**
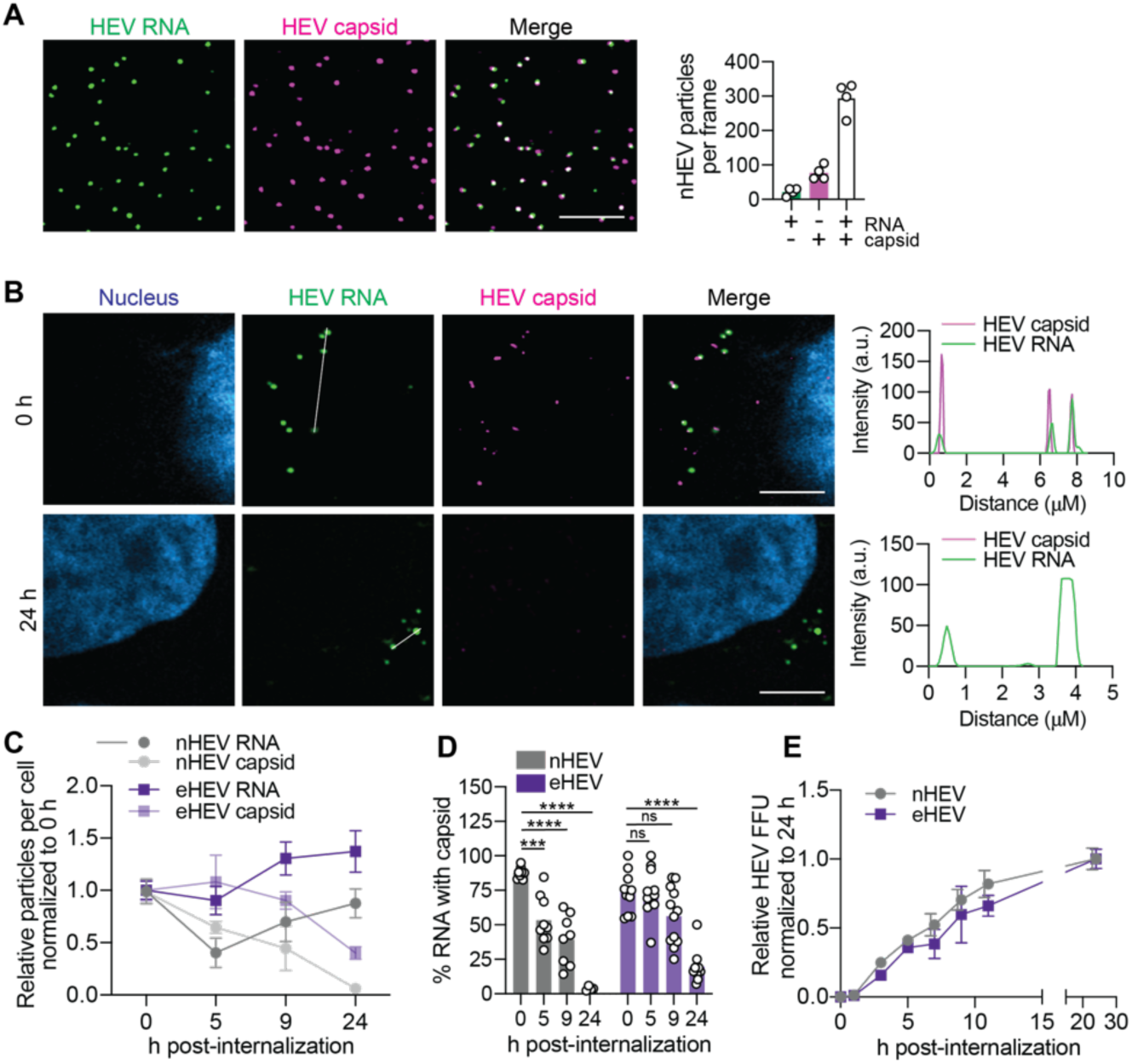
Co-detection of HEV capsid and genome allows analysis of the dynamics of nHEV and eHEV entry. (A) nHEV particles were immobilized on PEI-coated chamber slides and fixed. After permeabilization, capsids (magenta) were detected by immunofluorescence staining and HEV genomes (green) by RNA-FISH (version 1 kit) using the ORF1 probe. n = technical replicates from two independent virus productions. Scale bar = 10 μm. (B) S10-3 cells were inoculated with nHEV (MOI = 30 GE/cell) and incubated for 2 h at 4 °C to allow binding (upper image row) followed by inoculum removal and 24 h at 37°C for internalization (lower image row). After fixation, cells were stained for HEV capsids (magenta) and HEV genomes (green) detected by RNA-FISH (version 1 kit) using the ORF1 probe. Line graphs on the right show the fluorescence intensities of capsids and genomes measured across the region of interest indicated by the white line in the images shown on the left. Images are representatives of n = 6. Scale bar = 5 μm. (C) S10-3 cells were inoculated with nHEV (MOI = 30 GE/cell) or eHEV particles (MOI = 20 GE/cell) and allowed to bind for 2 h or 6 h, respectively, at 4 °C followed either by direct fixation (0 h) or internalization at 37 °C and fixation at indicated time points. HEV capsids and genomes were then detected by immunofluorescence staining and RNA-FISH (version 1 kit), respectively, and quantified using CellProfiler. The absolute numbers of detected nHEV or eHEV genomes and capsids over 24 h were normalized to 0 h. n = biological replicates from three independent experiments. (D) Images taken for the analysis in (C) were used to quantify the number of HEV genomes colocalising with capsid. Shown are the calculated percentages of HEV genomes with capsids out of the total number of detected genomes per cell. n = biological replicates from three independent experiments. (E) S10-3 cells were incubated with nHEV (MOI = 0.1 GE/cell) or eHEV particles (MOI = 5 GE/cell) on ice for 2 h and 6 h, respectively, followed by internalization at 37 °C. Ammonium chloride (NH_4_Cl) was added at different hours post-internalization to block viral entry and was replenished until the cells were fixed and analysed for infectivity by quantifying ORF2-positive FFUs 5 days post-infection. Images of entire infected wells were taken with a Zeiss CellDiscoverer 7 microscope and the number of FFU was counted manually. n = biological replicates from three independent experiments. For panels (A) to (D), the images were taken on a Zeiss Airyscan LSM900 confocal microscope. Images shown in (A) and (B) are maximum projections of 4 slices (thickness = 0.5 μm) and 2 slices (0.25 μm) respectively. Maximum projections of full z-series with a thickness of 10 μm were used for quantification of RNA and capsid particles with CellProfiler.

Next, we used this method to study the kinetics of HEV capsid uncoating (Fig. 3B). For nHEV, the number of capsids detected decreased steadily over the observation period. After 24 h, almost no capsids were detected anymore. In contrast, the number of HEV genomes remained relatively stable over the observation period. For eHEV, the number of capsids detected also decreased steadily. However, compared to nHEV, eHEV capsids decreased at a slower rate and only to 40% even 24 h after internalisation.

We then examined the percentage of HEV genomes that colocalised with the capsid over the course of 24 h post-internalisation. As shown in Fig. 3C, half of the incoming nHEV RNA was no longer colocalised with the capsid by 5 h post-internalisation, and this decreased to 5% by 24 h. In contrast, eHEV uncoating appeared to be much slower, with more than 50% of eHEV genomes still colocalised with the capsid at 9 h post-internalisation (Fig. 3C). After 24 h, we detected almost no HEV genomes colocalising with the capsid for nHEV particles and about 20% for eHEV particles. Since only the number of capsids, but not the genomes, decreased over the observation period (Fig. 3B), we concluded that HEV uncoating had occurred.

We further confirmed these results by time-of-addition experiments with ammonium chloride (NH_4_Cl), a weak lysosomotropic base known to promptly inhibit endocytosis. S10-3 cells were incubated with nHEV or eHEV on ice to allow particle binding. After removal of the inoculum, the cells were transferred to 37 °C to allow particle internalisation. We then added NH_4_Cl at indicated time points post-internalisation to block entry of particles that had not yet entered, in order to describe cell entry kinetics in a time-resolved manner. By counting FFUs 5 days post-infection, we found that only 50% of nHEV and 40% of HEV particles had completed their entry into cells at 7 h post-internalisation (Fig. 3E). This result confirmed our observations with the entry assay in Fig. 3C,D, suggesting that the cell entry processes of nHEV and HEV are rather slow compared to many other viruses. In particular, eHEV entry was slower and less productive than that of nHEV.

### Both eHEV and nHEV enter through the endocytic pathway but take differential routes

Upon ligand binding, integrins can be endocytosed in a clathrin-dependent manner and either degraded or recycled back to the plasma membrane^27^. Therefore, we hypothesized that nHEV interacts with ITGB1 upon binding and is internalised with ITGB1 by endocytosis. First, we used the endosomal acidification inhibitors bafilomycin A (BFA) and concanamycin A (Con A) and applied them to S10-3 cells prior to eHEV and nHEV infection (Fig. 4 A, B). Both treatments resulted in a dose-dependent decrease in eHEV and nHEV FFU numbers 5 days post-infection (Fig. 4A & B), in the absence of any drug-induced cell toxicity (Suppl. Fig. 6A). We verified that the drug treatments did not affect HEV replication (Suppl. Fig. 6B) and further confirmed their effect on nHEV infection in the hepatoma cell line HepG2.C3A as well as in primary human hepatocytes (Suppl. Figs. 6C & D).

**Figure 4:**
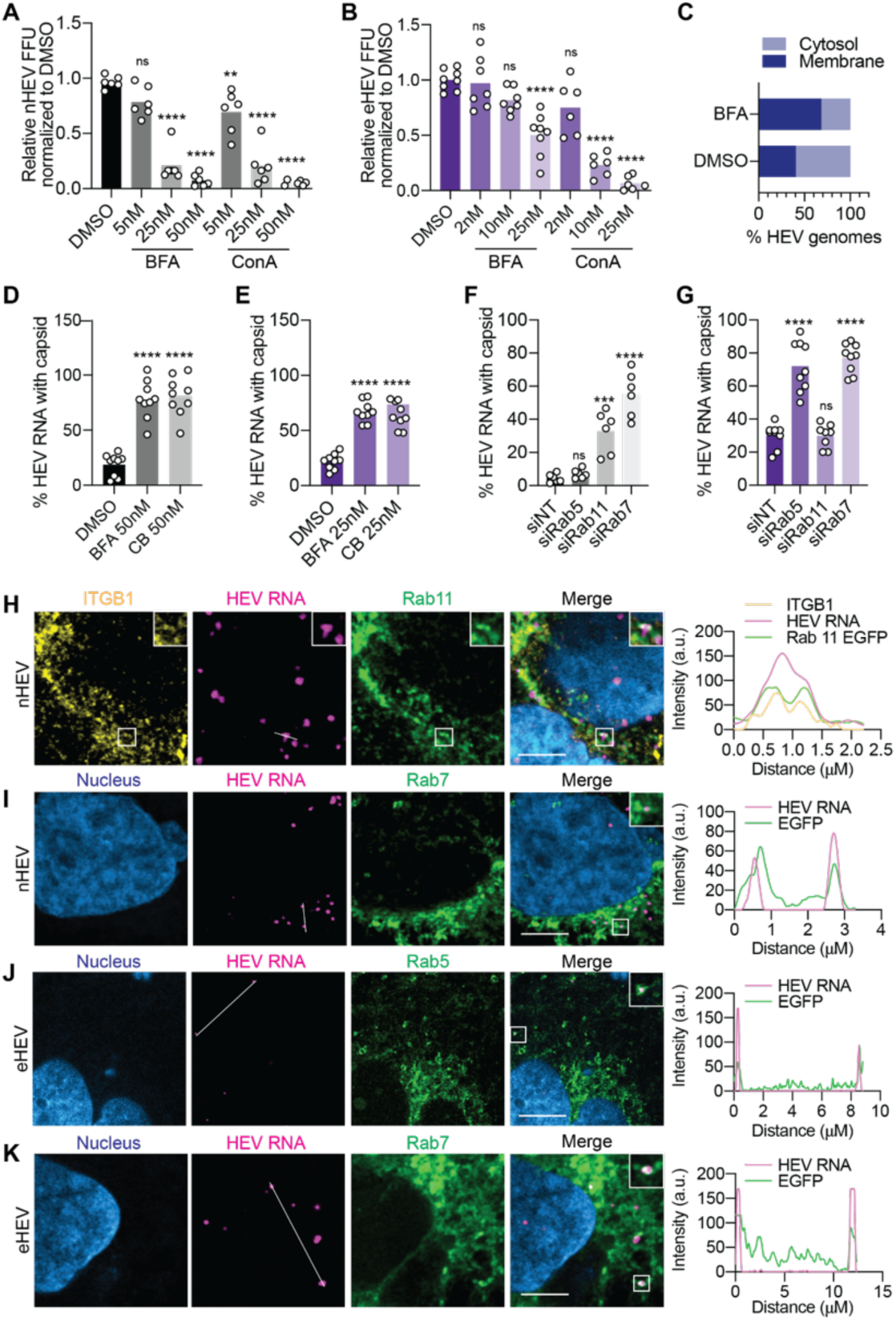
nHEV and eHEV particles traffic along different endocytic pathways. (A) and (B) S10-3 cells were treated with indicated concentrations of bafilomyin A (BFA), concanamyin A (ConA) or DMSO (mock) for 30 min prior to infecting with (A) nHEV (MOI = 0.1 GE/cell) or (B) eHEV (MOI = 5 GE/cell). Drugs and virus were removed after 24 hours and HEV infection was quantified by counting ORF2-positive FFUs 5 days post-infection. Images of entire infected wells were taken with a Zeiss CellDiscoverer 7 microscope and the number of FFU was counted manually. n = biological replicates from three independent experiments. (C) S10-3 cells were treated with 50 nM of BFA or DMSO before inoculation with nHEV (MOI = 20 GE/cell). The inoculum was removed 8 h later and replaced with fresh media containing the drug. 24 h later, cells were harvested followed by cell fractionation to extract membrane and cytosol fractions. HEV genome copies in each fraction were quantified by RT-qPCR. Shown are percentages of HEV genomes in each fraction in BFA and DMSO treated cells. N = 1. (D) and (E) S10-3 cells were treated with BFA, ConA or DMSO (mock) for 30 min before inoculation with (D) nHEV (MOI = 30 GE/cell) and (E) eHEV (MOI = 20 GE/cell). The inoculum was removed 8 h later and replaced with fresh media containing the drugs. 24 h later, the cells were fixed and HEV capsid and genomes detected by immunofluorescence staining and RNA-FISH (version 1 kit) using the ORF1 probe, respectively. Both capsids and genomes were quantified using CellProfiler. Shown are the percentages of HEV genomes co-localising with HEV capsids out of the total number of detected genomes per cell. n = biological replicates from three independent experiments. (F) and (G) S10-3 cells were transfected with 100 nM on-target pool siRNAs directed against Rab5, Rab7, Rab11, and a NT-control. 48 h post-transfection, the cells were inoculated with (F) nHEV (MOI = 30 GE/cell) or (G) eHEV (MOI = 20 GE/cell) and incubated for 8 h at 37 °C. 24 h post-inoculation, HEV capsid and HEV genomes were detected by immunofluorescence staining and RNA-FISH (version 1 kit) using the ORF1 probe, respectively. For (C) - (F), HEV genomes and capsids were quantified using CellProfiler. Shown are the calculated percentages of HEV genomes associated with capsids out of total number of detected genomes per cell. n = biological replicates from three independent experiments. (H) and (I) S10-3 cells ectopically expressing (H) Rab11-EGFP or (I) Rab7-EGFP were inoculated with nHEV particles (MOI = 30 GE/cell) and incubated for 15 min or 2 h at 37° C, respectively. The cells were fixed and stained against ITGB1 (yellow). HEV genomes (magenta) were detected by RNA-FISH (version 2 kit) using the ORF1 probe. Line graphs on the right show the fluorescence intensities of HEV genomes and GFP measured across the region of interest indicated by the white line in the images shown on the left. Images are representatives of n = 3. Scale bar = 5 μm. (J) and (K) S10-3 cells ectopically expressing Rab5-EGFP (J) or Rab7-EGFP (K) were inoculated with eHEV particles (MOI = 20 GE/cell) and incubated for 1 h or 8 h at 37° C, respectively. The cells were fixed and HEV genomes (magenta) were detected by RNA-FISH (version 2 kit) using the ORF1 probe. Line graphs on the right show the fluorescence intensities of HEV genomes and GFP measured across the region of interest indicated by the white line in the images shown on the left. Images are representatives of n = 3. Scale bar = 5 μm. (A) - (G) Statistical analysis was performed by one-way ANOVA **: *p* <0.01; ***: *p*<0.001; ****: p<0.0001; n.s., non-significant. Statistical comparisons for siRNA- or drug-treated groups were performed to the respective controls. For panels (D) to (K), the images were taken on a Zeiss Airyscan LSM900 confocal microscope. Images shown in (H) - (K) are maximum projections of 4 slices (thickness = 0.5 μm). Maximum projections of full z-series with thickness of 10 μm were used for quantification of HEV genomes and capsids with CellProfiler in (C) - (G).

Successful entry of the virus into the cell leads to release of the genome into the cytoplasm. We separated membranes from the cytosol after nHEV infection of BFA-treated and untreated S10-3 cells and found that the treatment led to an enrichment of detected HEV genomes in the membrane fraction. We also corroborated our findings using our entry assay based on HEV genome detection and capsid staining. We found a significant increase in capsid-associated genome-positive particles for both nHEV and eHEV upon endosomal inhibitor treatment compared to DMSO-treated cells (Fig. 4 E, F), suggesting that uncoating did not occur in the presence of the inhibitors.

Next, to investigate the specific endosomal compartments through which eHEV and nHEV particles are trafficked, we used siRNAs to knock down Rab5, Rab7 and Rab11, which are markers for the early, late, and recycling endosomes, respectively (Suppl. Fig. 7). We found that knockdown of Rab11, but not Rab5, resulted in a significant increase in capsid-associated genome nHEV particles (Fig. 4F). In contrast, knockdown of Rab5 but not Rab11 resulted in a significant increase in capsid-associated genome eHEV particles (Fig. 4G). Interestingly, Rab7 knockdown affected both eHEV and nHEV uncoating (Fig. 4F, G). We further confirmed the colocalisation of nHEV particles with Rab11 (Fig. 4H) and Rab7 (Fig. 4I) as well as the colocalisation of eHEV particles with Rab5 (Fig. 4J) and Rab7 (Fig. 4K) using high-resolution confocal microscopy.

Taken together, our results suggest that both nHEV and eHEV particles are highly dependent on the endocytic machinery for cell entry. Interestingly, we were also able to find ITGB1 and nHEV together in Rab11-positive endosomes 15 min after internalisation (Fig. 4E). This observation supports our hypothesis that the interaction with ITGB1 directs nHEV but not eHEV particles through Rab11-positive vesicles, with Rab11 being a hallmark of recycling endosomes.

### HEV uncoating in the lysosome requires cathepsin activity

Finally, we investigated whether eHEV and nHEV traffic through the final destination of endocytic cargoes, the lysosome. Since HEV particles are enterically transmitted and exposed to a highly acidic pH in the gut, we hypothesised that additional triggers are required for HEV uncoating. Lysosomal cathepsins have been shown to be involved in the entry of many viruses^28–31^. Therefore, we speculated that capsid processing by cathepsins might be important for HEV uncoating and subsequent genome release. As initial evidence, we found nHEV and eHEV particles in cellular vesicles positive for the lysosomal marker LAMP1 (Fig. 5A). We then treated cells with the cathepsin inhibitor E64 either before or after nHEV and eHEV infection and quantified HEV FFU 5 days post-infection. We found that the E64 inhibitor significantly reduced nHEV and eHEV infection when applied during the first 24 h of infection, which we further confirmed in PHHs (Suppl. Fig. 6D). In contrast, the application at 24 h post-infection had no effect on either eHEV or nHEV infection (Fig. 5B, C).

**Figure 5:**
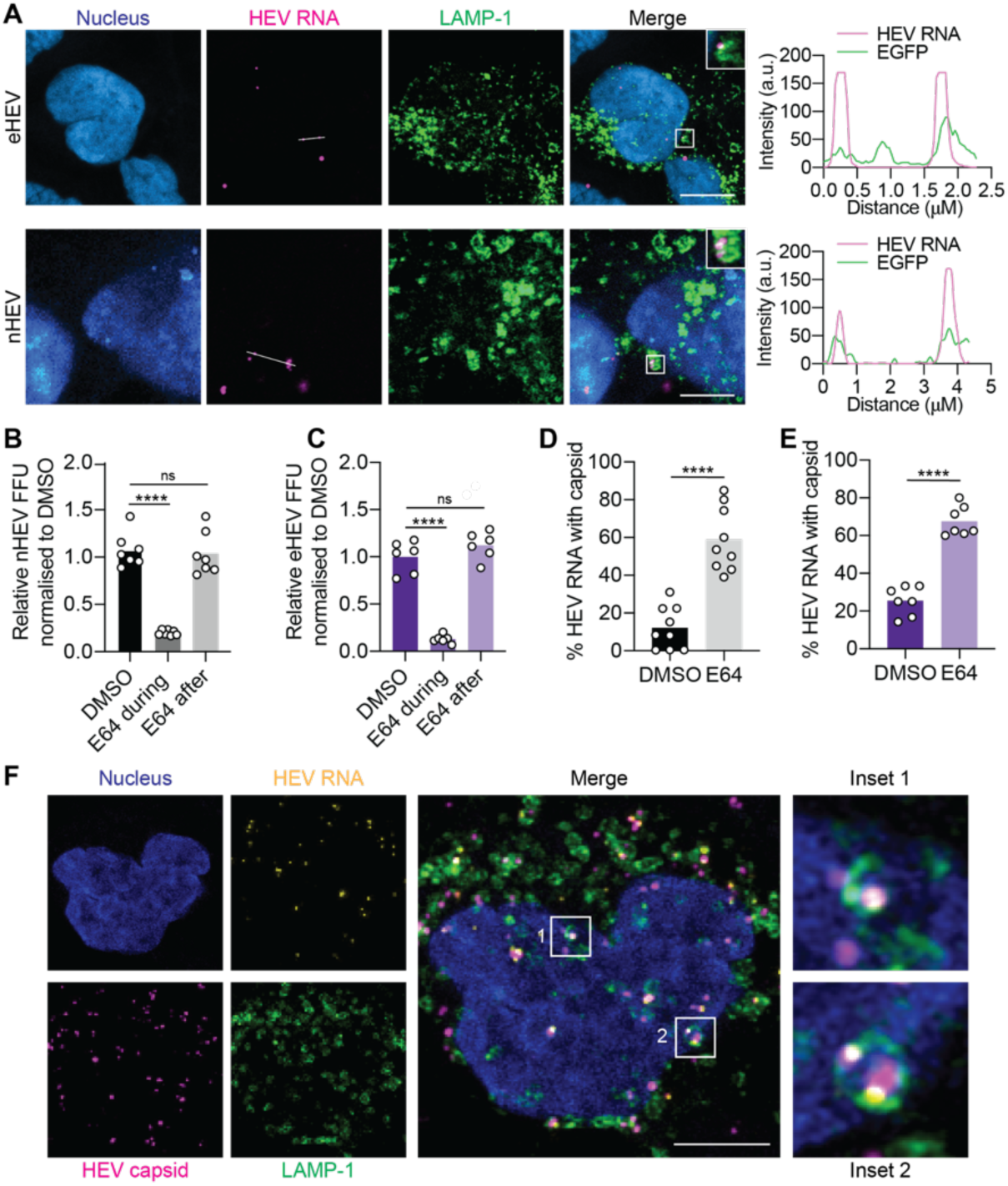
nHEV and eHEV particles require lysosomal cathepsin activity for cell entry. (A) S10-3 cells ectopically expressing LAMP-1-GFP were inoculated with eHEV (MOI = 20 GE/cell) or nHEV (MOI = 30 GE/cell) and incubated at 37 °C for 10 h or 7 h, respectively. The cells were fixed and HEV genomes (magenta) were detected by RNA-FISH using the ORF1 probe. Line graphs on the right show the fluorescence intensities of HEV genomes and GFP measured across the region of interest indicated by the white line in the images shown on the left. Images are representative of n = 3. Scale bar = 5 μm. (B) and (C) S10-3 cells were treated with 25 μM of cathepsin inhibitor E64 or DMSO (mock) for 30 min prior to infection with (B) nHEV (MOI = 30 GE/cell) or (C) eHEV (MOI = 20 GE/cell). Drugs and virus inoculum were removed 24 h later and HEV infection was quantified by counting ORF2-positive FFUs 5 days post-infection. Images of entire infected wells were taken with a Zeiss CellDiscoverer 7 microscope and the number of FFU was counted manually. n = biological replicates from three independent experiments. Statistical analysis was performed by one-way ANOVA **: *p* <0.01; ***: *p*<0.001; ****: p<0.0001; n.s., non-significant. (D) and (E) S10-3 cells were treated with 25 μM of E64 or DMSO (mock) for 30 min before inoculation with (D) nHEV (MOI = 30 GE/cell) or (E) eHEV (MOI = 20 GE/cell). The inoculum was removed 8 h later and replaced with fresh media containing the drugs. 24 h post-inoculation, cells were fixed and HEV capsid and genomes detected by immunofluorescence staining and RNA-FISH using the ORF1 probe, respectively. HEV genomes and capsids were quantified using CellProfiler. Shown are the calculated percentages of RNA particles associated with capsids out of the total number of detected genomes per cell. n = biological replicates from three independent experiments. Statistical analysis was performed by unpaired two-tailed Student’s t test. **: *p* <0.01; ***: *p*<0.001; ****: *p*<0.0001; n.s., non-significant. (F) S10-3 cells ectopically expressing LAMP-1-GFP were treated with 25 μM E64 and inoculated with nHEV (MOI = 30 GE/cell). 24 h post-inoculation, cells were fixed and HEV capsids (magenta) and HEV genomes (yellow) were detected by immunofluorescence staining and RNA-FISH, respectively. Images are representative of n = 3. For panels (A), (D), (E) and (F), the images were taken on a Zeiss Airyscan LSM900 confocal microscope. Images shown in (A) and (H) are maximum projections of 4 slices (thickness = 0.5 μm). Maximum projections of full z-series with thickness of 10 μm were used for quantification of HEV genomes and capsids with CellProfiler in (D) and (E).

Next, we confirmed the effect of E64 on nHEV and eHEV entry using our RNA-FISH based entry assay. We found a significant increase in capsid-associated genomes of both nHEV and eHEV particles upon E64 treatment compared to DMSO-treated cells (Fig. 5D, E). Furthermore, we observed entrapment of HEV capsid and colocalisation with the genome in LAMP1-positive lysosomes 24 h post-infection when treated with E64 (Fig. 5F), suggesting unsuccessful uncoating upon cathepsin inhibition. Taken together, these data suggest that both nHEV and eHEV particles uncoat in the lysosome and may require lysosomal cathepsins for genome release.

## Discussion

The early stages of the HEV life cycle are poorly understood. One obvious reason for this is the lack of suitable methods to study them separately from the later steps of the life cycle. Here, we used an RNA-FISH-based assay in combination with high-content imaging to study and describe authentic nHEV and eHEV cell entry steps, from the interaction with potential surface receptors to trafficking through the endocytic pathway. We provide experimental evidence that ITGB1 acts as a co-factor for nHEV but not eHEV entry. We further found that the two particle forms are differentially trafficked along the endocytic pathway and that lysosomal cathepsin activity is critical for particle uncoating of both forms.

### Integrin beta 1 is a co-host factor of nHEV entry

Previously, ITGA3 has been proposed as a co-host cell entry factor, but its beta partner has not yet been identified. As ITGA3 is thought to form a functional heterodimer with ITGB1 only, we sought to investigate its role in HEV cell entry. ITGB1 has been shown to be an entry factor for many viruses, including HAV^11^, rabies virus^21^, vaccinia virus^22^, and others. ITGB1 can bind to many alpha integrins and is ubiquitously expressed on various cell types. While the HEV capsid lacks the classic ITGB1 ligand motif, we have identified a reverse DGR motif in the protruding P-domain of ORF2 that can potentially mediate binding to ITGB1, albeit with low affinity^32^.

The expression of integrin heterodimers on the cell surface is tightly and dynamically regulated due to their critical cellular functions. When one subunit of integrins is downregulated or impaired, other subunits are known to compensate for their functions^33^. For example, knockdown of ITGB1 in various human and mouse breast cancer cell lines or kidney cells results in activation of integrin αvβ3^34^. This compensatory mechanism may explain the relatively mild phenotype we observed in ITGB1 knockout cells (Fig. 2). Notably, herpes simplex virus 1 (HSV-1) has been reported to use different integrin subunits as interchangeable receptors^35^. This ability allows the virus to adapt to alternative pathways in different cell types, thereby broadening its range of cellular targets for infection.

Indeed, the interaction with a promiscuous factor such as ITGB1 could explain the broad cell and species tropism of the HEV strain used in this study (reviewed in^36^). As the present work was limited to the use of this zoonotic strain, future studies should also include strains with a narrower tropism, such as the HEV-1 and -2 genotypes that are restricted to human infection. While ITGB1 is ubiquitously expressed, the alpha integrins seem to be more tissue specific, e.g. α3 and α6^37^ in epithelial cells and α10 in chondrocytes^38^ (reviewed in^16^). It would therefore be interesting to investigate whether ITGB1 heterodimerises with specific alphas that could be critical in mediating HEV entry into different tissues, such as the brain^39^ or intestine^40^, which have been described as permissive for nHEV infection in vitro and as potential reservoirs for HEV infection in chronic patients (reviewed in^41^).

### nHEV trafficking through the recycling endosome

In contrast to many other cell surface receptors that undergo synchronised ligand-induced internalisation and degradation, integrins are constantly recycled in cells and the recycling and internalisation routes of ITGB1 appear to be complex and highly regulated^27^. Following ligand binding, integrins can be internalised through clathrin-mediated endocytosis (CME) or clathrin-independent endocytosis (CIE), including caveolin-dependent pathways, micropinocytosis and clathrin-independent carriers (CLICs) (reviewed in^42^). There is evidence that some classes of beta-1 integrins are associated with Rab21^43^, which preferentially colocalises with caveolin-1 and regulates caveolin-mediated endocytic pathways^44^. In contrast, Rab5 mainly regulates clathrin-mediated endocytic pathways^45^. Once internalised, integrins are predominantly recycled back to the membrane mediated by Rab11 and the ADP-ribosylation factor 6 (ARF6), whereas integrin degradation is rather slow^45^. Our data suggest that the HEV capsid interacts directly or indirectly with ITGB1 and activates its internalisation possibly mediated by the recruitment of pFAK to Rab11+ recycling endosomes. Notably, many other viruses have been reported to enter cells through the recycling endosome^46–48^. It is possible that HEV is initially internalised into a Rab21-positive endosome and its trafficking through the recycling endosome is a by-product of the ITGB1 interaction. It would be interesting to investigate whether a fraction of nHEV particles is recycled back to the membrane and requires re-internalisation with ITGB1, which could explain the rather slow entry kinetics we observed (Fig. 3C-E). The HEV entry assay developed in this study could be used in the future to unravel the details of early nHEV entry.

Following internalisation, we found nHEV particles in LAMP1-positive compartments, which are likely to be endolysosomes. It is unusual for cargo in the recycling endosome to be targeted into the endolysosome for degradation. However, it has been shown that polyvalent ligands can cross-link receptor proteins that are normally recycled and that cross-linking of these receptors can alter their route to the degradation pathway^48,49^. For example, the non-enveloped canine parvovirus (CPV), has been shown to traffic through the recycling endosome but is detected in the lysosome 8 h post-internalisation^50^ and it has been proposed that redirection of CPV particles from the recycling pathway to the degradative pathway may be necessary for release of the virus from the vesicles.

Another possible link between the recycling endosome and the endolysosome is autophagosome formation. Rab11 has been proposed to regulate the fusion of MVBs with autophagosomes^51^, and LAMP1 is also known to be distributed among autophagic organelles^44^. In addition, Rab7 has been shown to play a key role in the autophagic pathway and the maturation of autophagosomes^52,53^ and there is evidence that some viruses, such as adenovirus and human echovirus 7, use the autophagy machinery for cell entry^54^. It is possible that the autophagy machinery could promote the recruitment of molecular motors (such as dynein) to the ruptured endosome and assist in the exit of the HEV genome from the endosomes^55^. Thus, it is possible that nHEV particles also use the autophagic pathway, since our data suggest that nHEV entry does not follow the classical degradative endocytic pathway to the lysosome via Rab5, but rather involves Rab11 (Fig. 4).

### Potential underlying mechanism of low infectivity of eHEV particles

In contrast to nHEV particles, eHEV particles, which are unlikely to contain virus-encoded proteins exposed on the quasi-envelope, do not appear to interact with ITGB1 (Fig. 2). A recent study has shown that the phosphatidylserine (PtdSer) receptor TIM-1 mediates eHEV entry through PtdSer embedded in the quasi-envelope. Similarly, PtdSer is displayed on the surface of eHAV and initial attachment of eHAV to cells is mediated in part by TIM-1^14^. In agreement with the study by Yin *et al*^9^, we found that eHEV binding to cells was less efficient than nHEV binding, probably due to the envelope which prevents specific uptake and internalisation via ITGB1.

In addition, however, we found that eHEV uncoating also appeared to be less efficient with roughly 25% of detected genomes still being associated with capsid, even after 24 h internalization (Fig. 3D). A possible explanation for this slower and or inefficient entry process of eHEV could be the premature fusion of the envelope with endosomal membranes. Studies have shown that eHEV acquires its membrane from the trans-Golgi network^56^ prior to budding into multivesicular bodies (MVBs)^57,58^. MVBs can fuse with late endosomes, where the cellular content in MVBs can either be packaged into lysosomes for degradation or released as exosomes^59^. Since eHEV has a membrane similar to that of an exosome, it is possible that eHEV particles fuse prematurely with endosomal membranes, leading to the release of intact capsids into the cytoplasm and thus to unproductive cell entry.

An additional rate-limiting determinant for eHEV cell entry could be the need to remove the quasi-envelope to allow exposure of the capsid to cathepsins. Unsuccessful uncoating in the lysosome could then lead to subsequent degradation, which may altogether explain the lower infectivity of eHEV compared to nHEV. Thus, while the quasi-envelope provides obvious benefits to HEV, such as non-cytolytic release and protection from neutralising antibodies, it also renders the particles much less infectious (Suppl. Fig. 4).

Many open questions remain about the cell entry pathways of both nHEV and eHEV particles. The slow entry kinetics of both particle forms remain intriguing. Many viruses, such as human immunodeficiency virus^60^, HSV^61^, or SARS-CoV2^62^ complete the entry process within a few hours, whereas nHEV and eHEV uncoating takes more than 12 h (Fig. 3D, E). Future studies should aim at further elucidating the molecular mechanism in a time-resolved manner of each step. In this manuscript, we have laid the groundwork and provide a novel entry assay to enable such studies. A better description of the HEV cell entry steps could not only lead to the development of therapeutic interventions, but also to a better understanding of HEV tropism and extrahepatic manifestations.

## Acknowledgments

The authors gratefully acknowledge Dr. Suzanne Emerson and Dr. Britta Brügger for sharing reagents and Andrew Freistaedter for excellent technical support. We acknowledge Dr. Vibor Laketa, head of the Infectious Diseases Imaging Platform (IDIP) at the University Hospital Heidelberg for expert support. pSpCas9(BB)-2A-GFP (PX458) was a gift from Feng Zhang (Addgene plasmid # 48138; http://n2t.net/addgene:48138; RRID:Addgene_48138) and pWPI was a gift from Didier Trono (Addgene plasmid # 12254; http://n2t.net/addgene:12254; RRID:Addgene_12254). We thank Ann-Kathrin Mehnert for valuable and helpful discussions.

## Author contributions

Conceptualization, R.M.F, S.L. and V.L.D.T.; Methodology, R.M.F, Z.E.; Investigation, R.M.F, Z.E., J.A.W, J.M., M.K., D.T., J.H.; Resources: E.S., P.Y.L; Software, R.M.F.; Data analysis, R.M.F. and V.L.D.T.; Writing-original draft, R.M.F. and V.L.D.T.; Final draft, R.M.F., P.Y.L., S.L., and V.L.D.T.; Supervision, V.L.D.T.; Funding, V.L.D.T.

## References

1 Horvatits, T., Schulze Zur Wiesch, J., Lutgehetmann, M., Lohse, A. W. & Pischke, S. The Clinical Perspective on Hepatitis E. Viruses 11, doi:10.3390/v11070617 (2019).

2 Nimgaonkar, I., Ding, Q., Schwartz, R. E. & Ploss, A. Hepatitis E virus: advances and challenges. Nature reviews. Gastroenterology & hepatology, doi:10.1038/nrgastro.2017.150 (2017).

3 Purdy, M. A. et al. ICTV Virus Taxonomy Profile: Hepeviridae. J Gen Virol 98, 2645–2646, doi:10.1099/jgv.0.000940 (2017).

4 Fu, R. M., Decker, C. C. & Dao Thi, V. L. Cell Culture Models for Hepatitis E Virus. Viruses 11, doi:10.3390/v11070608 (2019).

5 Meister, T. L., Bruening, J., Todt, D. & Steinmann, E. Cell culture systems for the study of hepatitis E virus. Antiviral research 163, 34–49, doi:10.1016/j.antiviral.2019.01.007 (2019).

6 Das, A. et al. Cell entry and release of quasi-enveloped human hepatitis viruses. Nat Rev Microbiol 21, 573–589, doi:10.1038/s41579-023-00889-z (2023).

7 Yin, X., Li, X. & Feng, Z. Role of Envelopment in the HEV Life Cycle. Viruses 8, doi:10.3390/v8080229 (2016).

8 Dao Thi, V. L., et al. Stem cell-derived polarized hepatocytes. Nat Commun 11, 1677, doi:10.1038/s41467-020-15337-2 (2020).

9 Yin, X., Ambardekar, C., Lu, Y. & Feng, Z. Distinct Entry Mechanisms for Nonenveloped and Quasi-Enveloped Hepatitis E Viruses. J Virol 90, 4232–4242, doi:10.1128/JVI.02804-15 (2016).

10 Takahashi, M. et al. Monoclonal antibodies raised against the ORF3 protein of hepatitis E virus (HEV) can capture HEV particles in culture supernatant and serum but not those in feces. Archives of virology 153, 1703–1713, doi:10.1007/s00705-008-0179-6 (2008).

11 Rivera-Serrano, E. E., Gonzalez-Lopez, O., Das, A. & Lemon, S. M. Cellular entry and uncoating of naked and quasi-enveloped human hepatoviruses. eLife 8, doi:10.7554/eLife.43983 (2019).

12 Yin, X. & Feng, Z. Hepatitis E Virus Entry. Viruses 11, doi:10.3390/v11100883 (2019).

13 Schrader, J. A. et al. EGF receptor modulates HEV entry in human hepatocytes. Hepatology 77, 2104–2117, doi:10.1097/HEP.0000000000000308 (2023).

14 Corneillie, L. et al. The phosphatidylserine receptor TIM1 promotes infection of enveloped hepatitis E virus. Cellular and molecular life sciences : CMLS 80, 326, doi:10.1007/s00018-023-04977-4 (2023).

15 Shiota, T. et al. Integrin α3 is involved in non-enveloped hepatitis E virus infection. Virology 536, 119–124, doi:10.1016/j.virol.2019.07.025 (2019).

16 Barczyk, M., Carracedo, S. & Gullberg, D. Integrins. Cell Tissue Res 339, 269–280, doi:10.1007/s00441-009-0834-6 (2010).

17 Das, A. et al. Gangliosides are essential endosomal receptors for quasi-enveloped and naked hepatitis A virus. Nat Microbiol 5, 1069-+, doi:10.1038/s41564-020-0727-8 (2020).

18 Holla, P., Ahmad, I., Ahmed, Z. & Jameel, S. Hepatitis E Virus Enters Liver Cells Through a Dynamin-2, Clathrin and Membrane Cholesterol-Dependent Pathway. Traffic 16, 398–416, doi:10.1111/tra.12260 (2015).

19 Zhou, Z., Xie, Y., Wu, C. & Nan, Y. The Hepatitis E Virus Open Reading Frame 2 Protein: Beyond Viral Capsid. Front Microbiol 12, 739124, doi:10.3389/fmicb.2021.739124 (2021).

20 Csernalabics, B., et al. Efficient formation and maintenance of humoral and CD4 T cell immunity targeting naked but not quasi-enveloped virions in acute-resolving hepatitis E infection. BioRxiv. doi: 2023/563038. 10.1101/2023.10.19.563038

21 Shuai, L. et al. Integrin β1 Promotes Peripheral Entry by Rabies Virus. Journal of Virology 94, doi:10.1128/Jvi.01819-19 (2020).

22 Izmailyan, R. et al. Integrin β1 Mediates Vaccinia Virus Entry through Activation of PI3K/Akt Signaling. Journal of Virology 86, 6677–6687, doi:10.1128/Jvi.06860-11 (2012).

23 Kaminsky, P. M., Keiser, N. W., Yan, Z. Y., Lei-Butters, D. C. M. & Engelhardt, J. F. Directing Integrin-linked Endocytosis of Recombinant AAV Enhances Productive FAK-dependent Transduction. Molecular Therapy 20, 972–983, doi:10.1038/mt.2011.295 (2012).

24 Gianni, T., Gatta, V. & Campadelli-Fiume, G. αβ-integrin routes herpes simplex virus to an entry pathway dependent on cholesterol-rich lipid rafts and dynamin2. P Natl Acad Sci USA 107, 22260–22265, doi:10.1073/pnas.1014923108 (2010).

25 Guan, J. L. Integrin Signaling Through FAK in the Regulation of Mammary Stem Cells and Breast Cancer. Iubmb Life 62, 268–276, doi:10.1002/iub.303 (2010).

26 Kumar, C. C. Signaling by integrin receptors. Oncogene 17, 1365–1373, doi:DOI 10.1038/sj.onc.1202172 (1998).

27 Arjonen, A., Alanko, J., Veltel, S. & Ivaska, J. Distinct Recycling of Active and Inactive β1 Integrins. Eur J Cancer 48, S116–S116, doi:Doi 10.1016/S0959-8049(12)71156-5 (2012).

28 Bar, S., Daeffler, L., Rommelaere, J. & Nuesch, J. P. Vesicular egress of non-enveloped lytic parvoviruses depends on gelsolin functioning. PLoS Pathog 4, e1000126, doi:10.1371/journal.ppat.1000126 (2008).

29 Ebert, D. H., Deussing, J., Peters, C. & Dermody, T. S. Cathepsin L and cathepsin B mediate reovirus disassembly in murine fibroblast cells. Journal of Biological Chemistry 277, 24609–24617, doi:10.1074/jbc.M201107200 (2002).

30 Zhao, M. M. et al. Cathepsin L plays a key role in SARS-CoV-2 infection in humans and humanized mice and is a promising target for new drug development. Signal Transduct Tar 6, doi:ARTN 134 10.1038/s41392-021-00558-8 (2021).

31 Qiu, Z. Z. et al. Endosomal proteolysis by cathepsins is necessary for murine coronavirus mouse hepatitis virus type 2 spike-mediated entry. Journal of Virology 80, 5768–5776, doi:10.1128/Jvi.00442-06 (2006).

32 Spitaleri, A. et al. Structural basis for the interaction of with the RGD-binding site of αvβ3 integrin. Journal of Biological Chemistry 283, 19757–19768, doi:10.1074/jbc.M710273200 (2008).

33 Samarzija, I. et al. Integrin Crosstalk Contributes to the Complexity of Signalling and Unpredictable Cancer Cell Fates. Cancers (Basel) 12, doi:10.3390/cancers12071910 (2020).

34 Hayashida, T., Jones, J. C. R., Lee, C. K. & Schnaper, H. W. Loss of β1-Integrin Enhances TGF-β1-induced Collagen Expression in Epithelial Cells via Increased αvβ3-Integrin and Rac1 Activity. Journal of Biological Chemistry 285, 30741–30751, doi:10.1074/jbc.M110.105700 (2010).

35 Gianni, T., Salvioli, S., Chesnokova, L. S., Hutt-Fletcher, L. M. & Campadelli-Fiume, G. αvβ6-and αvβ8-Integrins Serve As Interchangeable Receptors for HSV gH/gL to Promote Endocytosis and Activation of Membrane Fusion. Plos Pathogens 9, doi:ARTN e1003806 10.1371/journal.ppat.1003806 (2013).

36 Meng, X. J. Expanding Host Range and Cross-Species Infection of Hepatitis E Virus. Plos Pathogens 12, doi:ARTN e1005695 10.1371/journal.ppat.1005695 (2016).

37 Hierck, B. P. et al. Variants of the Alpha(6)Beta(1)-Laminin Receptor in Early Murine Development - Distribution, Molecular-Cloning and Chromosomal Localization of the Mouse Integrin Alpha(6)-Subunit. Cell Adhes Commun 1, 33–53, doi:Doi 10.3109/15419069309095680 (1993).

38 Varas, L. et al. α10 integrin expression is up-regulated on fibroblast growth factor-2-treated mesenchymal stem cells with improved chondrogenic differentiation potential. Stem Cells Dev 16, 965–978, doi:10.1089/scd.2007.0049 (2007).

39 Zhou, X. et al. Hepatitis E Virus Infects Neurons and Brains. Journal of Infectious Diseases 215, 1197–1206, doi:10.1093/infdis/jix079 (2017).

40 Marion, O. et al. Hepatitis E virus replication in human intestinal cells. Gut, doi:10.1136/gutjnl-2019-319004 (2019).

41 Kamar, N., Marion, O., Abravanel, F., Izopet, J. & Dalton, H. R. Extrahepatic manifestations of hepatitis E virus. Liver international : official journal of the International Association for the Study of the Liver 36, 467–472, doi:10.1111/liv.13037 (2016).

42 Moreno-Layseca, P., Icha, J., Hamidi, H. & Ivaska, J. Integrin trafficking in cells and tissues. Nature Cell Biology 21, 122–132, doi:10.1038/s41556-018-0223-z (2019).

43 Pellinen, T. et al. Small GTPase Rab21 regulates cell adhesion and controls endosomal traffic of β1-integrins. J Cell Biol 173, 767–780, doi:DOI 10.1083/jcb.200509019 (2006).

44 Shikanai, M. et al. Rab21 regulates caveolin-1-mediated endocytic trafficking to promote immature neurite pruning. Embo Rep 24, doi:10.15252/embr.202254701 (2023).

45 Caswell, P. T. & Norman, J. C. Integrin trafficking and the control of cell migration. Traffic 7, 14–21, doi:10.1111/j.1600-0854.2005.00362.x (2006).

46 Mannova, P. & Forstova, J. Mouse polyomavirus utilizes recycling endosomes for a traffic pathway independent of COPI vesicle transport. J Virol 77, 1672–1681, doi:10.1128/jvi.77.3.1672-1681.2003 (2003).

47 Greene, W. & Gao, S. J. Actin dynamics regulate multiple endosomal steps during Kaposi’s sarcoma-associated herpesvirus entry and trafficking in endothelial cells. PLoS Pathog 5, e1000512, doi:10.1371/journal.ppat.1000512 (2009).

48 Johns, H. L., Berryman, S., Monaghan, P., Belsham, G. J. & Jackson, T. A Dominant-Negative Mutant of rab5 Inhibits Infection of Cells by Foot-and-Mouth Disease Virus: Implications for Virus Entry. Journal of Virology 83, 6247–6256, doi:10.1128/Jvi.02460-08 (2009).

49 Marsh, E. W., Leopold, P. L., Jones, N. L. & Maxfield, F. R. Oligomerized Transferrin Receptors Are Selectively Retained by a Lumenal Sorting Signal in a Long-Lived Endocytic Recycling Compartment. J Cell Biol 129, 1509–1522, doi:DOI 10.1083/jcb.129.6.1509 (1995).

50 Suikkanen, S. et al. Role of recycling endosomes and lysosomes in dynein-dependent entry of canine parvovirus. Journal of Virology 76, 4401–4411, doi:10.1128/Jvi.76.9.4401-4411.2002 (2002).

51 Fader, C. M., Sánchez, D., Furlán, M. & Colombo, M. I. Induction of autophagy promotes fusion of multivesicular bodies with autophagic vacuoles in K562 cells. Traffic 9, 230–250, doi:10.1111/j.1600-0854.2007.00677.x (2008).

52 Gutierrez, M. G., Munafó, D. B., Berón, W. & Colombo, M. I. Rab7 is required for the normal progression of the autophagic pathway in mammalian cells. J Cell Sci 117, 2687–2697, doi:10.1242/jcs.01114 (2004).

53 Hyttinen, J. M. T., Niittykoski, M., Salminen, A. & Kaarniranta, K. Maturation of autophagosomes and endosomes: A key role for Rab7. Bba-Mol Cell Res 1833, 503–510, doi:10.1016/j.bbamcr.2012.11.018 (2013).

54 Montespan, C. et al. Multi-layered control of Galectin-8 mediated autophagy during adenovirus cell entry through a conserved PPxY motif in the viral capsid. Plos Pathogens 13, doi:ARTN e1006217 10.1371/journal.ppat.1006217 (2017).

55 Viret, C., Roziéres, A. & Faure, M. Autophagy during Early Virus-Host Cell Interactions. J Mol Biol 430, 1696–1713, doi:10.1016/j.jmb.2018.04.018 (2018).

56 Nagashima, S. et al. The membrane on the surface of hepatitis E virus particles is derived from the intracellular membrane and contains trans-Golgi network protein 2. Archives of virology 159, 979–991, doi:10.1007/s00705-013-1912-3 (2014).

57 Nagashima, S. et al. Characterization of the Quasi-Enveloped Hepatitis E Virus Particles Released by the Cellular Exosomal Pathway. Journal of Virology 91, doi:UNSP e00822-17 10.1128/JVI.00822-17 (2017).

58 Nagashima, S. et al. Hepatitis E virus egress depends on the exosomal pathway, with secretory exosomes derived from multivesicular bodies. Journal of General Virology 95, 2166–2175, doi:10.1099/vir.0.066910-0 (2014).

59 Scott, C. C., Vacca, F. & Gruenberg, J. Endosome maturation, transport and functions. Semin Cell Dev Biol 31, 2–10, doi:10.1016/j.semcdb.2014.03.034 (2014).

60 Gallo, S. A. et al. Kinetic studies of HIV-1 and HIV-2 envelope glycoprotein-mediated fusion. Retrovirology 3, doi:Artn 90 10.1186/1742-4690-3-90 (2006).

61 McClain, D. S. & Fuller, A. O. Cell-specific kinetics and efficiency of herpes simplex virus type 1 entry are determined by two distinct phases of attachment. Virology 198, 690–702, doi:10.1006/viro.1994.1081 (1994).

62 Koch, J. et al. TMPRSS2 expression dictates the entry route used by SARS-CoV-2 to infect host cells. Embo Journal 40, doi:ARTN e107821 10.15252/embj.2021107821 (2021).

