## Supplementary Material for "A high-content RNA-based imaging assay reveals integrin beta 1 as a cofactor for cell entry of non-enveloped hepatitis E virus"

**Supplementary Material Fu et al.**

**Supplementary Methods**

*Primary human hepatocytes*

Primary human hepatocytes (PHH) were purchased from PRIMACYST(Lot #: CHM2225-HE-Z) and seeded onto collagen-coated 24-well plate (PRIMACYST) following the manufacturer's protocol precisely. In brief, frozen vials of PHH were thawed in thawing media (PRIMACYST) and $4.1x10^5$ cells per well were seeded in plating medium (PRIMACYST) and allowed to attach for 6 h at 37 ℃ and 5% CO2. After cells were attached, the media was removed and cells were washed twice with warm PBS and replenished with culture medium (PRIMACYST). 24 h post-seeding, the cells were treated with the indicated inhibitors and infected with nHEV at MOI = 0.5 GE/cell.

*Renilla luciferase replication assay*

S10-3 cells were electroporated with the RNA encoding a subgenomic HEV replicon[1] and seeded onto 12-well plates. Cell culture supernatant was harvested every 24 h and passed through a 0.45 µm Millex HV filter (Millipore), and subsequently stored at -80 °C. Fresh medium was then added to the culture, and the incubation process was resumed. Upon completing the experiment, all collected media were thawed at the same time, and 20 µl from each sample was subjected to a luciferase activity assay using the Renilla luciferase assay system (Promega). 50 µl of culture medium was added to individual wells of a black flat-bottom 96-well plate (Corning). Subsequently, 1 nM Coelenterazine was diluted in PBS and 60 µl per well was added. The emission was measured 10 s after the addition of the substrate with a Centro LB 963 Microplate luminometer (Berthold).

*Quantitative (real-time) reverse transcription-PCR*

RNA was extracted from cell lysate or purified virus with Trizol reagent (Sigma) following the manufacturer's recommended protocol. cDNA was synthesized using the iScript cDNA Synthesis Kit (BioRad). HEV RNA GEs were quantified using the SYBR Green Master mix (BioRad) and primers targeting the HEV genome[2] (Fw: GGTGGTTTCTGGGGTGAC, Rv: AGGGGTTGGTTGGATGAA). RPS11 gene was used as a housekeeping control (Fw: GCCGAGACTATCTGCACTAC , Rv: ATGTCCAGCCTCAGAACTTC). The qPCR reaction was carried out in a BioRad CFX Maestro instrument.

*Transcriptome analysis*

To investigate integrin expression levels in primary human hepatocytes (PHH), previously published datasets were mined[3,4] (GEO accession numbers GSE140591 and GSE135619). Transcriptomic analyses were performed using CLC Genomics Workbench (QIAGEN). Raw FASTQ files were mapped against the hg19 human reference genomes with annotated gene and mRNA tracks. Gene expression was calculated for individual transcripts as RPKM.

### Supplementary Figures

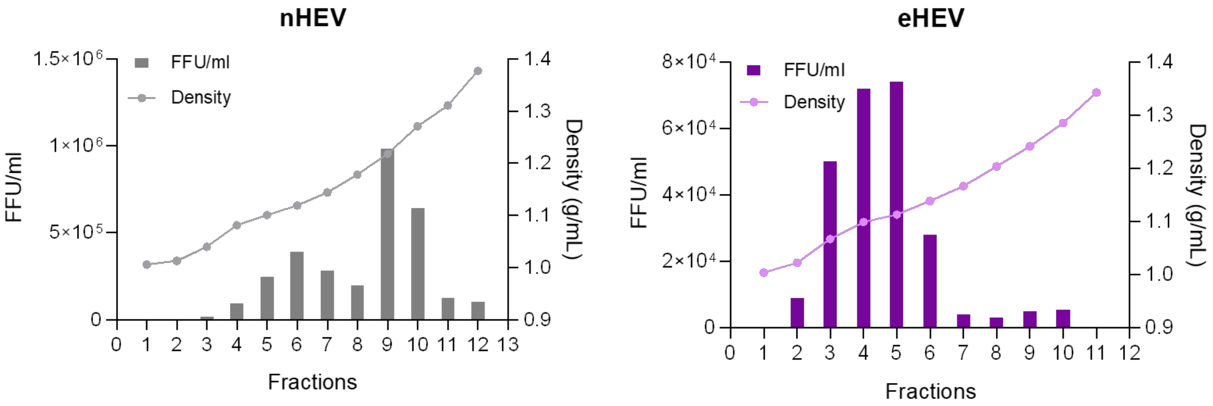


**Supplementary Figure 1. Characterization of nHEV and eHEV particles.**
Buoyant density of HEV particles in cell lysates (left panel) or released into supernatant fluids (right panel) of S10-3 cells electroporated with HEV RNA purified through isopycnic iodixanol gradient centrifugation. HEV RNA in fractions was determined by qRT-PCR. HEV infectivity was determined by infecting S10-3 cells and quantifying FFU 5 days post-infection. Images of entire infected wells were taken with a Zeiss CellDiscoverer 7 microscope and the number of FFU was counted manually. N = 1 biological replicate.

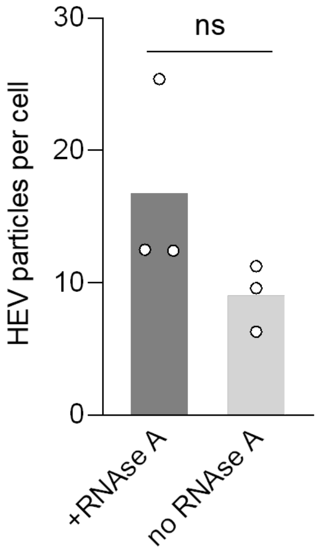


**Supplementary Figure 2. Pre-treatment of HEV with RNase A does not affect HEV genome detection via RNAscope.**
nHEV particles were pre-incubated with RNAse A at 25 µg/ml for 1 h at 37°C. S10-3 cells were then inoculated with treated or untreated virus (MOI = 10) for 6 h. Cells were fixed and HEV genomes were detected by RNA-FISH (version 1 kit) using the ORF1 probe. The images were taken on a Leica SP8 confocal microscope. Maximum projections of full z-series with a thickness of 10 µm were used for quantification of HEV genomes. The detected HEV genomes were quantified using CellProfiler. HEV particles per cell were calculated by dividing the total number of detected HEV genomes by the number of nuclei in an image frame. n = biological replicates from one experiment. Statistical analysis was performed by unpaired two-tailed Student's t test. **: $p < 0.01$; ***: $p<0.001$; ****: $p<0.0001$; n.s., non-significant.

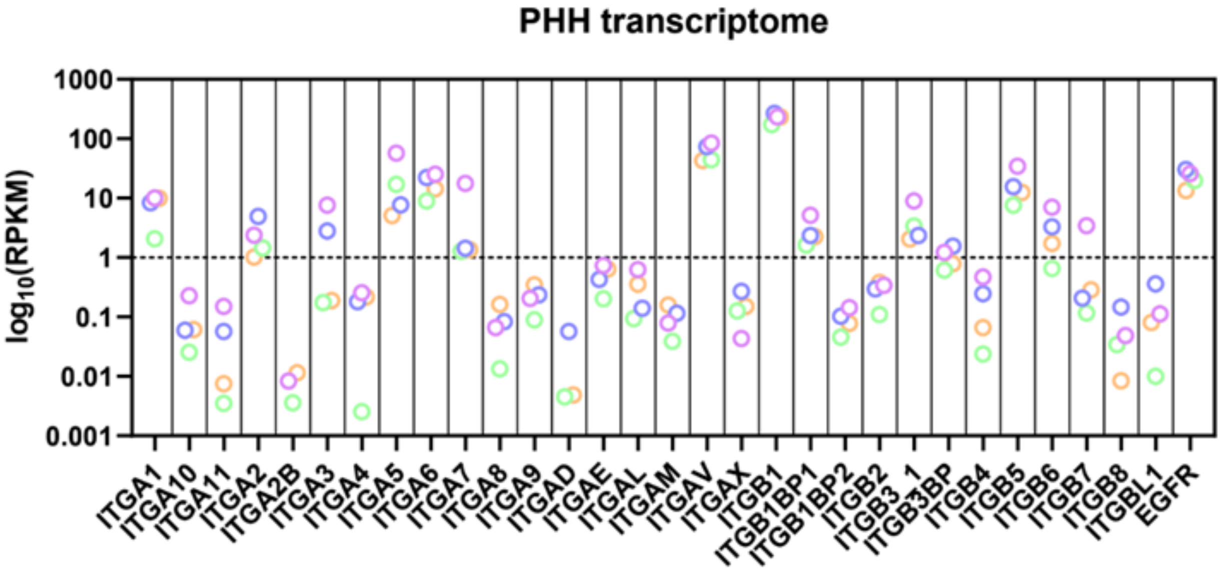


**Supplementary Figure 3**. **Integrin expression levels in primary human hepatocytes (PHH).**
RNAseq data showing expression levels of selected α and β integrins in PHH expressed in reads per kilobase of transcript (RPKM).

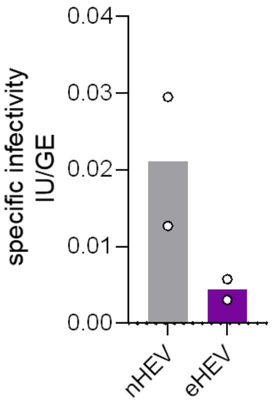


**Supplementary Figure 4. Specific infectivity of nHEV and eHEV.**
Specific infectivity of nHEV and eHEV represented as infectious units per genome copy (IU/GE). HEV RNA was extracted from gradient-purified virus and quantified by qPCR. HEV infectivity was determined by infecting S10-3 cells and quantifying FFU 5 days post-infection. Images of entire infected wells were taken with a Zeiss CellDiscoverer 7 microscope and the number of FFU was counted manually. n = biological replicate from two independent virus productions.

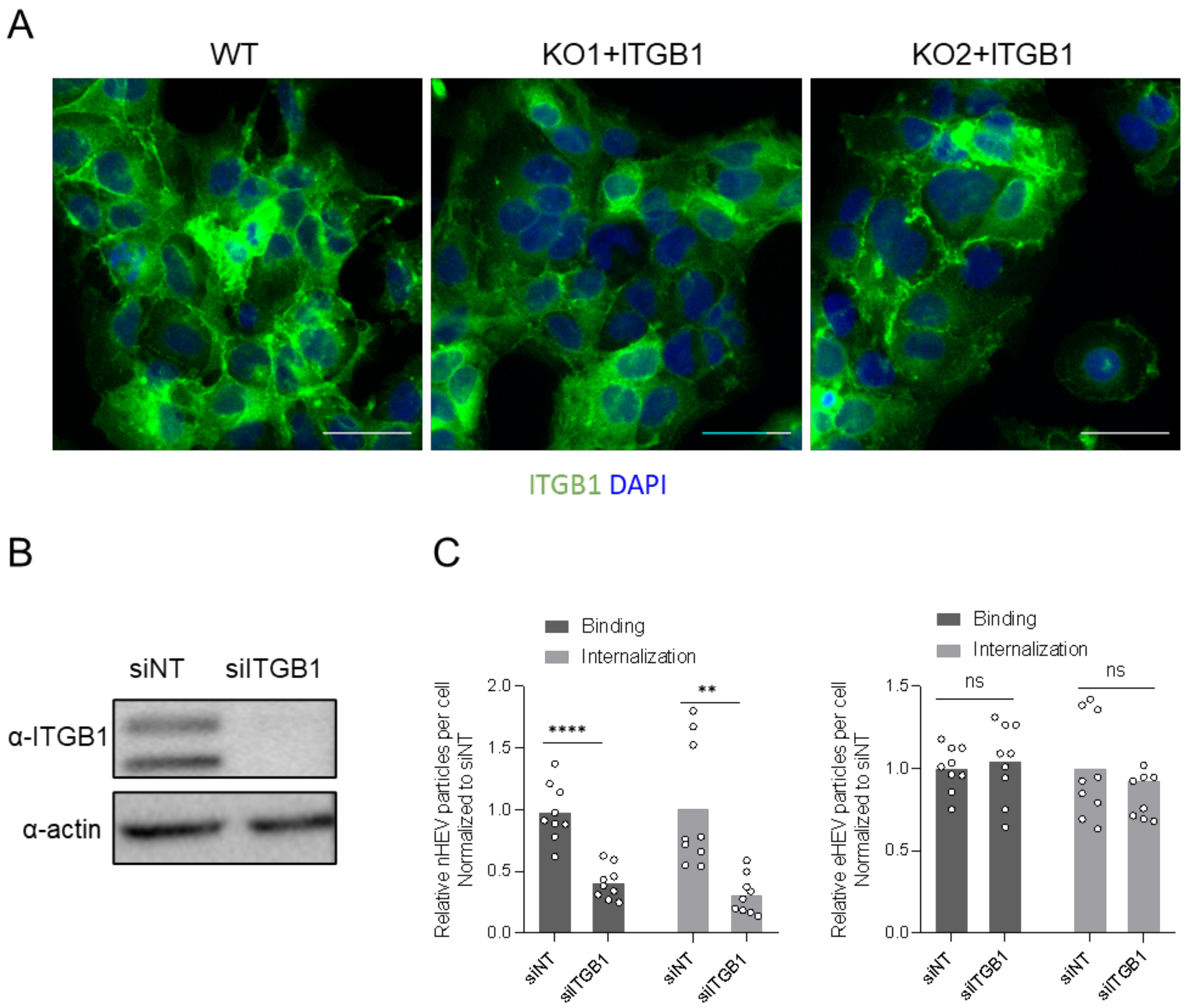


**Supplementary Figure 5. Rescue of ITGB1 in KO cells and effect of ITGB1 knockdown in HEV entry in S10-3 cells.**

(A) Ectopic ITGB1 expression in S10-3 ITGB1 KO cells. S10-3 ITGB1 KO cells were transduced with lentiviruses expressing ITGB1 and selected with puromycin. S10-3 WT and KO cells with ectopic ITGB1 were fixed and stained with DAPI (blue, nucleus) and against ITGB1 (green). The images were acquired on a Zeiss CellDiscoverer 7 microscope. Scale bar = 50 µm. (B) Western blot analysis of lysates harvested from S10-3 cells 48 h post-transfection with 100 nM on-target pool siRNAs directed against the ITGB1 gene (siITGB1) or a non-target control (siNT). (C) siITGB- and siNT-transfected S10-3 cells were inoculated with nHEV (left panel) (MOI = 30 GE/cell) or eHEV particles (right panel) (MOI = 20 GE/cell), 48 h post-transfection. The cells were incubated for 2 h or 6 h, respectively, at 4 °C to allow particle binding. Cells were then either fixed or the inoculum removed and shifted to incubation at 37 °C for 6 h to allow HEV particle internalization. The images were taken on a Zeiss Airyscan LSM900 confocal microscope. Maximum projections of full z-series with a thickness of 10 µm were used for quantification of RNA and capsid particles with CellProfiler. n = biological replicates from three independent experiments. Statistical analysis was performed by one-way ANOVA **: *p* <0.01; ****: p<0.0001; ns, non-significant.

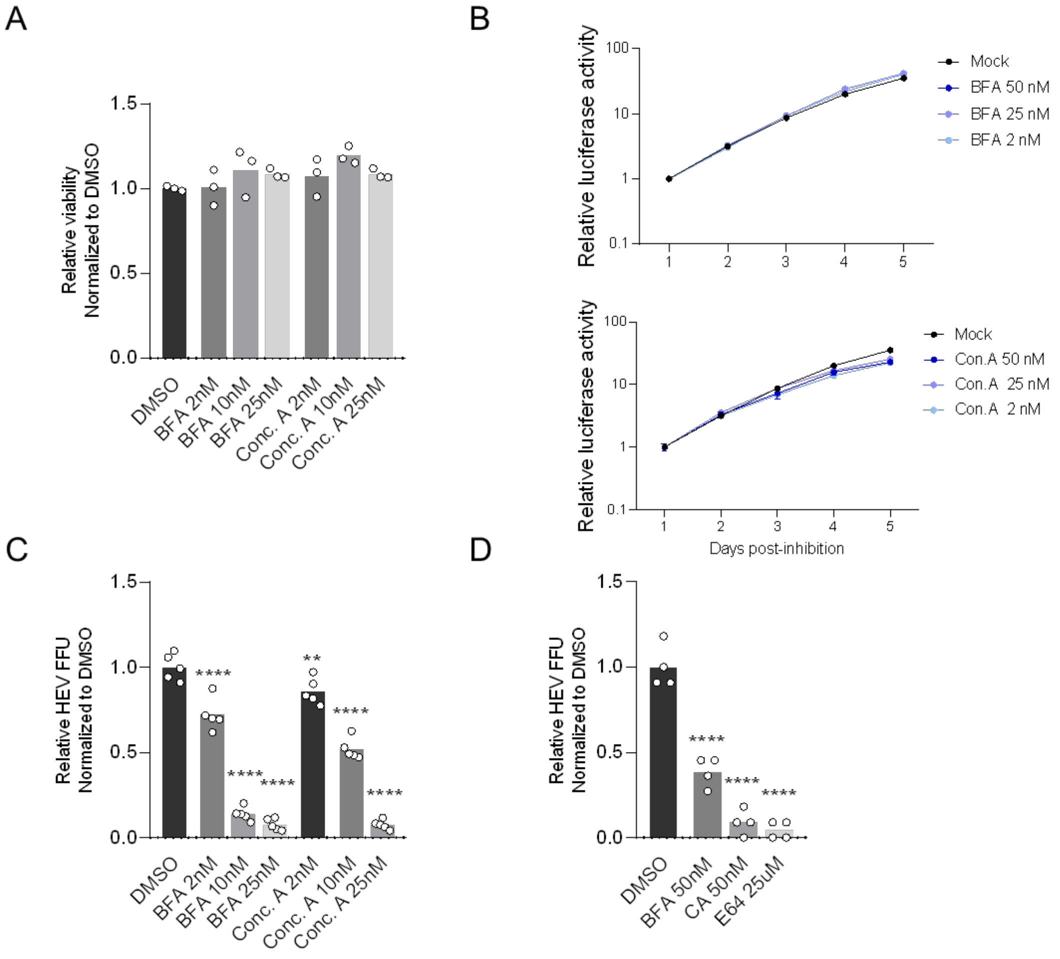


**Supplementary Figure 6. Effect of endocytic inhibitors on cell viability, viral replication, and HEV infection in HepG2/C3A and PHH.**

(A) S10-3 cells were treated with indicated concentrations of bafilomyin A (BFA), concanamyin A (ConA) or DMSO for 18h and cell viability was measured by an MTT assay. n = biological replicates from one experiment. (B) S10-3 cells were electroporated with an HEV-GLuc subgenomic replicon and treated with the inhibitors in (A). HEV replication was quantified by measuring luciferase activity in the supernatant during days 1 to 4 post-electroporation. n = biological replicates from two independent experiments. (C) HepG2/C3A cells were treated with indicated concentrations of bafilomyin A (BFA), concanamyin A (ConA) or DMSO (mock) for 30 min prior to infecting with nHEV (MOI = 0.1 GE/cell). Drugs and virus were removed after 24 h and HEV infection was quantified by counting ORF2-positive FFUs 5 days post-infection. Images of entire infected wells were taken with a Zeiss CellDiscoverer 7 microscope and the number of FFU was counted manually. n = biological replicates from three independent experiments. (D) Primary human hepatocytes (PHH) were treated with indicated concentrations of BFA, ConA and E64 in the same manner as in (C) and inoculated with nHEV (MOI = 0.5 GE/cell). FFUs were quantified 3 days post-infection. Images of entire infected wells were taken with a Zeiss CellDiscoverer 7 microscope and the number of FFU was counted manually. n = biological replicates from one experiment. Statistical analysis was performed by one-way ANOVA **: $p$ <0.01; ****: p<0.0001.

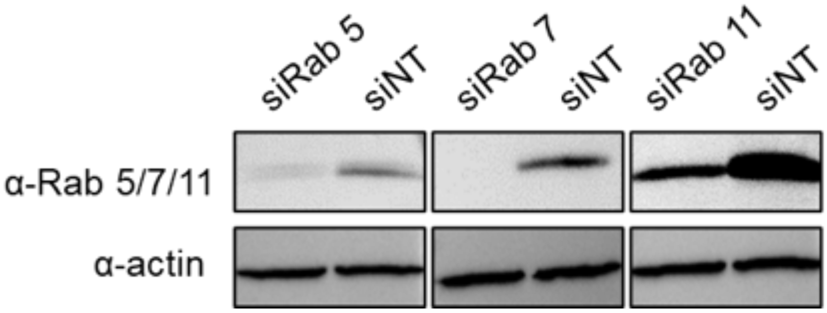


**Supplementary Figure 7. siRNA-mediated knockdown of Rab5, Rab7 and Rab11**.
Western blot analysis of lysates harvested from S10-3 cells 48 h post-transfection with 100 nM on-target pool siRNAs directed against the ITGB1 gene (siITGB1) or a non-target control (siNT).

### TO Supplementary References


1 Shukla, P. *et al.* Adaptation of a genotype 3 hepatitis E virus to efficient growth in cell culture depends on an inserted human gene segment acquired by recombination. *J Virol* **86**, 5697-5707, doi:10.1128/JVI.00146-12 (2012).

2 Jothikumar, N., Cromeans, T. L., Robertson, B. H., Meng, X. J. & Hill, V. R. A broadly reactive one-step real-time RT-PCR assay for rapid and sensitive detection of hepatitis E virus. *Journal of virological methods* **131**, 65-71, doi:10.1016/j.jviromet.2005.07.004 (2006).

3 Brown, R. J. P. *et al.* Liver-expressed and limit hepatitis C virus cross-species transmission to mice. *Science advances* **6**, doi:ARTN eabd323 10.1126/sciadv.abd3233 (2020).

4 Todt, D. *et al.* Robust hepatitis E virus infection and transcriptional response in human hepatocytes. *P Natl Acad Sci USA* **117**, 1731-1741, doi:10.1073/pnas.1912307117 (2020).
